# Modelling capture efficiency of single cell RNA-sequencing data improves inference of transcriptome-wide burst kinetics

**DOI:** 10.1101/2023.03.06.531327

**Authors:** Wenhao Tang, Andreas Christ Sølvsten Jørgensen, Samuel Marguerat, Philipp Thomas, Vahid Shahrezaei

## Abstract

Gene expression is characterised by stochastic bursts of transcription that occur at brief and random periods of promoter activity. The kinetics of gene expression burstiness differs across the genome and is dependent on the promoter sequence, among other factors. Single-cell RNA sequencing (scRNA-seq) has made it possible to quantify the cell-to-cell variability in transcription at a global genome-wide level. However, scRNA-seq data is prone to technical variability, including low and variable capture efficiency of transcripts from individual cells. Here, we propose a novel mathematical theory for the observed variability in scRNA-seq data. Our method captures burst kinetics and variability in both cell size and capture efficiency, which allows us to propose several likelihood-based and simulation-based methods for the inference of burst kinetics from scRNA-seq data. Using both synthetic and real data, we show that the simulation-based methods provide an accurate, robust and flexible tool for inferring burst kinetics from scRNA-seq data. In particular, in supervised manner, a simulation-based inference method based on neural networks proves to be accurate and useful in application to both allele and non-allele specific scRNA-seq data.

## 1 Introduction

Gene expression is stochastic in nature due to the random timing of chemical reactions involving low numbers of key molecular players, such as genes and mRNAs, as well as the coupling to other variable cellular processes, such as the cell cycle. This stochasticity gives rise to cell-to-cell phenotypic variability in a population of genetically identical cells, with a broad impact on cellular function.

Over the last twenty years, a considerable body of research combining experimental and mathematical studies has provided a deep understanding of the sources and consequences of this kind of biomolecular noise [1, 2, 3]. Single-cell imaging studies of fluorescently tagged proteins were the first to quantify gene expression noise [4]. Pioneering experimental and mathematical research broadly classified the sources of stochastic gene expression as either intrinsic due to random timing of the reactions involved in gene expression or as extrinsic due to the fluctuations of other relevant cellular factors [5]. Also, direct time-lapse imaging and inference from snap-shot data revealed gene expression could occur in bursts [6, 7, 8, 9, 10]. Methods such as the single-molecule Fluorescence In Situ Hybridisation (smFISH) and MSN2 system allowed for the quantification of gene expression noise and burstiness at the mRNA level [11, 12]. Most recently, the development of single-cell RNA sequencing (scRNA-seq) has made it possible to map global transcript counts in many cells and many genes routinely and cheaply [13]. scRNA-seq data can reveal biophysical mechanisms of gene regulation when they are combined with mechanistic models [14, 15]. However, due to additional technical variability in scRNA-seq data, inferring burst kinetics from such data is a challenging mathematical and statistical problem [13].

As mRNA copy numbers are typically low, it is generally well accepted that transcription is dominated by intrinsic noise [7] but the cell cycle can contribute to extrinsic expression noise [16]. Recent work has shown that transcription is coupled to cell size in eukaryotic systems, which underlies mRNA concentration homeostasis and also underlies extrinsic variability in gene expression [17, 18, 19, 20]. Accounting for cell size and cellular context transcription is reported to be nonbursty following a Poisson distribution in some cellular systems [17, 21]. However, more generally transcription is observed to be bursty and is modelled well using a so-called telegraph model, in which transcription switches between on and off states [7]. The telegraph model is analysed theoretically extensively and it is known that it admits Beta-Poisson distribution at steady-state [22, 23, 7, 24, 25]. At the bursty limit of transcription the solution of the telegraph model can be approximated as a negative-binomial distribution characterised by burst size and burst frequency [7, 24, 26, 27, 28]. Moreover, the negative binomial (NB) distribution is a versatile over-dispersed distribution that is commonly used in bulk and scRNA-seq studies to model gene expression capturing both biological and technical dispersion [29, 30, 31, 32].

The inference of parameters of mathematical models of stochastic gene expression from single-cell data is an important and challenging problem. Depending on the type of model, type of data, and the form of extrinsic noise, a range of different approaches have been developed recently to tackle this kind of inference problem [17, 33, 34, 35, 36, 37, 38, 39, 40, 41]. The inference of gene expression burst kinetics from scRNA-seq data has its own unique challenges due to specific kind of technical variability, complexity and sparsity of such data. Several recent studies have used single allele-specific scRNAs-seq data to map global burst kinetics genome-wide based on the Beta-Poisson distribution solution of the telegraph model [42, 43, 25, 44]. However, it is still an open question how to take into account the extrinsic biological and technical variability such as variation in cell size and capture efficiency in such methods [45]. The model by [44] considers the cell-specific variations via spike-ins data, which is an experimental control that is not commonly available. In addition, the model by [44] does not properly account for low and variable capture rates in scRNA-seq protocols. Meanwhile, the recent work by [42] applies Maximum Likelihood Estimation (MLE) directly on the raw scRNA-seq counts, hereby ignoring the cell-specific extrinsic variations. Ignoring extrinsic noise in such inference can inflate the amount of variability attributed to intrinsic noise and could lead to misleading estimates of the burst kinetics.

Here, we revisit the problem of statistical inference of the parameters of gene expression from scRNA-seq data focusing on the role of extrinsic variability. We present a mathematical model of gene expression measured by scRNA-seq. Our model appropriately accounts for the extrinsic variability introduced by cell-to-cell variations in scRNA-seq capture efficiency and cell size. To estimate the gene-specific kinetic parameters, we implement and compare four different inference schemes: MLE, methods of moments estimation (MME), an Approximate Bayesian Computation (ABC) rejection sampling algorithm, and using direct likelihood free inference based on a neural network (NN) implementation [46]. We benchmark these inference methods in a series of applications to synthetic and real data and discuss which methods work best.

## 2 Methods

### 2.1 Theory and model

The classic model for stochastic gene expression is the so-called telegraph model (Fig. 1a). It is known that the Chemical Master Equation of the telegraph model results in a Beta-Poisson distribution for the mRNA at steady state [22, 7, 47].

**Figure 1:**
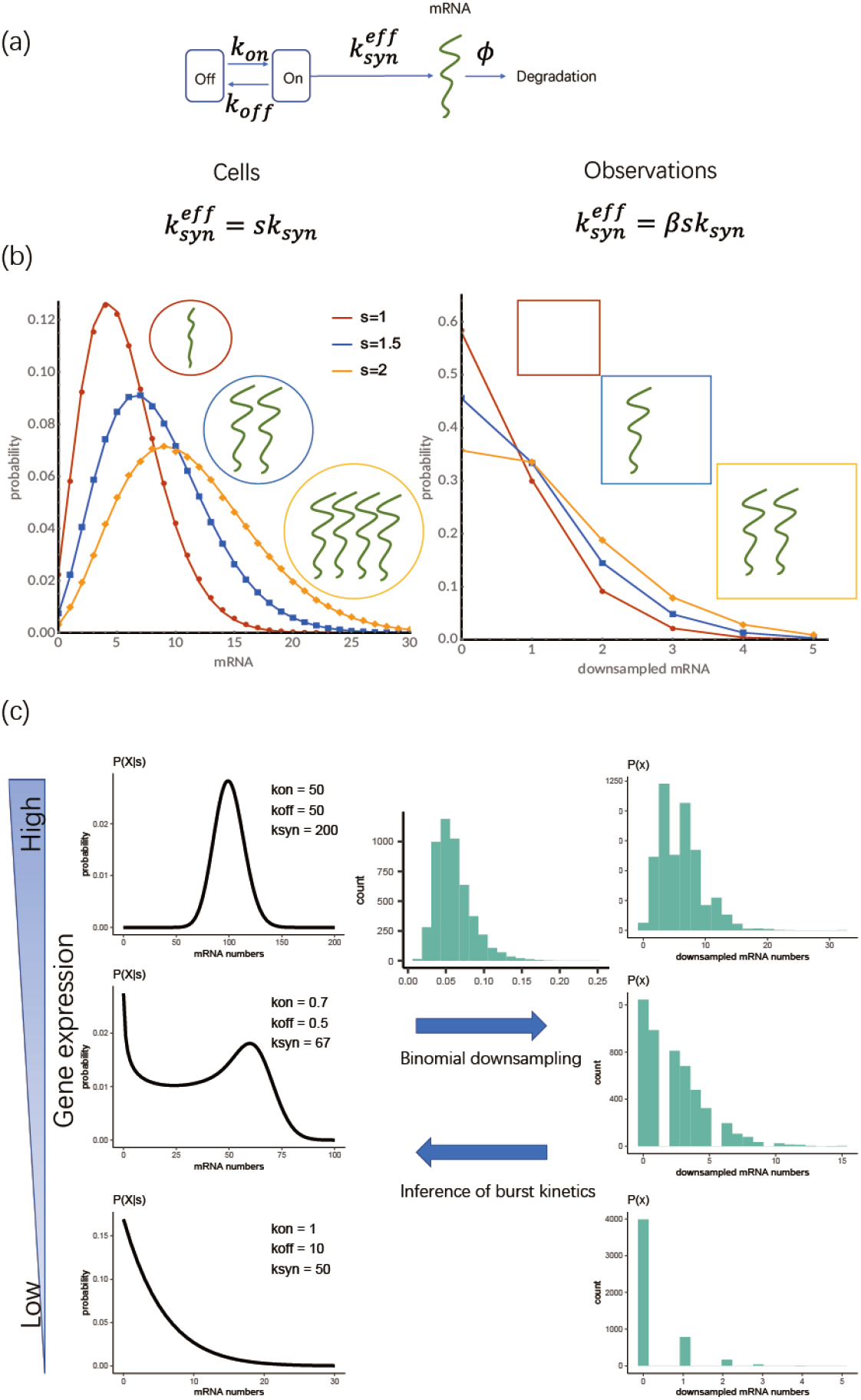
Model of stochastic gene expression and the effect of cell size and sequencing capture efficiency on observed transcript count distributions. **(a)** An illustration of the telegraph model of stochastic gene expression and its associated parameters. Gene switches between an inactive and active state and mRNAs are transcribed only from the active state. **(b)** Illustration of downsampling in scRNA-seq with a constant *β* = 0.5 (note that in reality *β* tends to be smaller and varies across the cells. Effective transcription rate 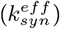 is proportional to cell size in original transcript counts (right) and both cell size and capture efficiency in the observed counts (right). **(c)** Distributions of original mRNA counts in cells with constant size for three specific parameters sets for the telegraph model (left) and their corresponding downsampled distribution (right). Distribution of cell specific capture efficiencies (*β*) used in downsampling is illustrated in the middle top arrow (sampled from a log-normal distribution as described in Section 2.4.3). The challenge is to use the downsampled observed count distribution that is also affected by variability in capture efficiency and cell size to infer the parameters of the original distribution (middle bottom arrow).

This result is only valid in the absence of any extrinsic noise and cell cycle effects with a gene with a constant transcription rate (*k*_*syn*_). However, as discussed in the introduction, gene expression is coupled to cell size and is, therefore, affected by the cell cycle [17, 21]. Moreover, we have recently shown that the telegraph model satisfies the so-called stochastic concentration homeostasis condition when the transcription rate scales with cell size (*s*) [48]. This notion implies that the transcript counts (*X*_*ij*_) of gene *i* in cell *j* in a population of growing and dividing cells (Fig. 1) is distributed as follows:

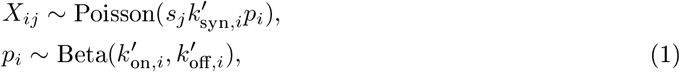

where *s*_*j*_ is the cell size, and 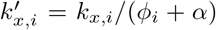 denote the gene-specific synthesis and promoter switching rates scaled by the effective degradation rate. The latter comprises the gene specific degradation rate *φ*_*i*_ and the exponential growth rate *α* of the population.

During scRNA-seq, only a fraction of transcripts in each cell is captured. As we have recently, the transcript counts observed in scRNA-seq data can be well modelled by a binomial model with a cell-specific capture efficiency (probability) denoted by *β*_*j*_ [31]. Intuitively, the binomial model is a natural choice as each transcript in a given cell is captured with the same cell-specific probability *β*_*j*_. Notably, the binomial model can explain the statistics of drop-out events without a need to invoke any zero-inflation models [31, 32].

Using this binomial model, one can show that the distribution of observed transcripts (*x*_*ij*_) in a cell of size *s*_*j*_ and capture efficiency *β*_*j*_ still follows the Beta-Poisson distribution but with a scaled effective synthesis rate:

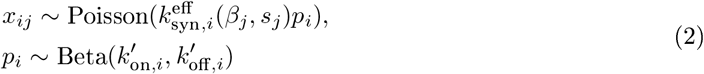

with 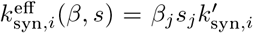 denoting an effective transcription rate for the observed counts. The observed counts *x* are necessarily lower than the actual original counts *X* and we therefore also refer to these as the downsampled counts. The dependence of the actual and observed transcript distributions on *β* and *s* is illustrated in Fig. 1b and c. This distribution then represents the correct likelihood function that should be used in the inference of kinetic rates from scRNA-seq data as it takes the biological variability introduced by the cell size and technical variability introduced by the capture efficiency into account. In the following, the kinetic rates of the model are defined relative to the effective decay rate, and as we are dealing with snap-shot data (and assuming steady state), we will omit the primes on the scaled rates.

### 2.2 Estimation of the capture efficiency

Let *x*_*ij*_ (*i* ∈ {1, 2, …, *P*} and *j* ∈ {1, 2, …, *Q*}) denote the number of transcripts reported for the *i*^th^ gene in the *j*^th^ cell in a scRNA-seq study. The measurements could be single allele or non-single allele, depending on the protocol. We introduce 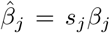 for cell *j*, which can simply be estimated using an appropriate cell-specific scaling factor [13] from the raw data (*x*_*ij*_). One simple such measure of the cell-specific scale factor is the total number of counts:

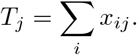

We posit that 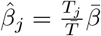 where 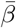 is the average capture efficiency in the scRNA-seq protocol and 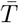 is the average total count across cells. It can be estimated, for example, from smFISH data [31, 49]. Throughout most of this paper, 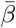 is set to 0.06 in accordance with Klein et al. [50].

### 2.3 Likelihood- and moment-based estimation of burst kinetics

#### 2.3.1 Maximum Likelihood Estimation (MLE)

Let dBP denote the Beta-Poisson distribution. A parameter estimation method proposed in the literature ([42]) is obtained by the maximising the log-likelihood:

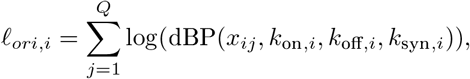

where *x*_*ij*_ are the raw single-allele scRNA-seq counts for gene *i* in cell *j* as above, and the sum runs over all cells in the data set - here, we assume the observations are independent. This approach attributes all variations in the transcript counts to the intrinsic stochastic properties of gene expression (in this case, in the framework of the telegraph model). As this method does not use any normalisation and relies on raw scRNA-seq counts, we have termed it in this study as bare MLE (denoted as BMLE).

There are two ways to evaluate *ℓ*_*ori*_ numerically. First, one might employ the so-called integral method used in previous studies (see [44] and [42] for more details). Alternatively, one can use the analytic form derived by Raj et al. [7] and Amrhein et al. [27]:

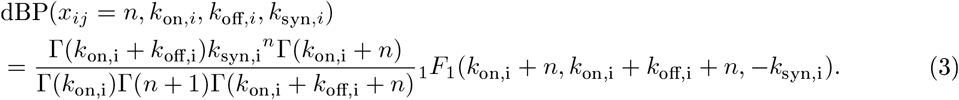

The limitation of this method is that it implicitly assumes perfect capture efficiency. This could in principle be overcome by considering the log-likelihood function derived from model (2):

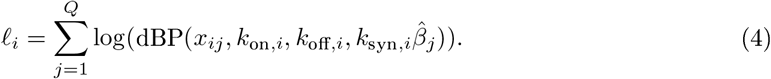

This method is denoted simply as MLE in this study (in contrast to BMLE). The integral method is not suitable for our modified MLE method, however, as we consider cell-specific parameters which would require us to marginalise over all cells. Since it is not possible to efficiently vectorize this approach, the computation becomes very time-consuming. We hence utilised the analytic form (3) to take cell-specific parameters into consideration. However, this method is numerically challenging since it involves evaluation of Eq. (3) for various expression levels and parameter combinations. This circumstance can lead to uncontrolled numerical errors that impair the accuracy of the MLE (Fig. S1).

#### 2.3.2 Method of Moments Estimation (MME)

As an alternative to the MLE method, we consider the Method of Moments Estimation as presented by Larsson et al. [42]. In this study, we denote this approach as the bare MME (BMME) as it is based on raw counts. The method considers the first three moments 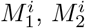, and 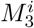 of each gene that can be estimated based on the raw single-allele scRNA-seq data (cf. [42]):

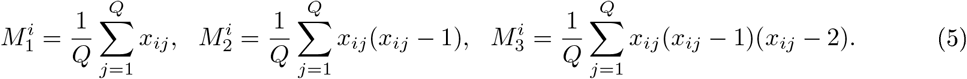

For brevity of notation, we will omit the index *i* from the moments as the parameters of each gene can be estimated independently. One can then define the quantities

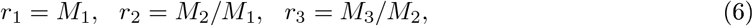

that are related to the kinetic parameters:

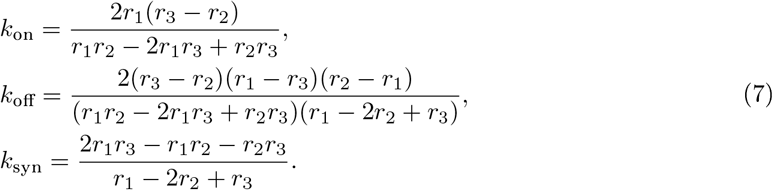

To take variations in the capture efficiency 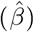 into consideration, we propose a heuristic approach that uses 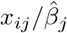 instead of the raw *x*_*ij*_ in the estimation of the moments. We denote this method simply as MME in contrast to BMME.

### 2.4 Simulation-based estimation of burst kinetics

#### 2.4.1 Approximate Bayesian Computation (ABC)

When dealing with our modified MLE method, there are numerical instability issues, and the optimisation is challenging. Meanwhile, it is known that MME may lead to biased estimates. Alternative methods that avoid these issues are likelihood-free approaches that imply sampling simulations. In our case, we sample from our analytical distribution (Eq. (4)).

Here, we employ ABC rejection sampling using priors that are constructed based on Section 2.4.3. In ABC, one relies on a distance measure between the data and the simulations. For this purpose, we employ the Hellinger distance since it has good properties with respect to model misspecification [51].

The data, one gene’s expression 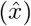 are a vector of scRNA-seq counts of length *Q* cells associated with another vector of capture efficiencies 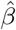. For each gene, the algorithm goes through the following steps:

1. We draw kinetic parameter sets from the prior: ***θ***^**⋆**^ *∼ π*(***θ***) (Section 2.4.3). Some parameter sets are filtered out to make sure that *M*_1_ ∈ {*M*_5%_, *M*_95%_}, where *M*_5%_ and *M*_95%_ indicate the 5^th^ and 95^th^ percentiles of l,000 *M*_1_ estimates from bootstrapped MME (that is, we sample cells with replacement 1,000 times and hence obtain l,000 *M*_1_ estimates for each gene).
2. We simulate data (*x*^**⋆**^) by sampling a vector of gene expressions from 2 using ***θ***^**⋆**^ and a cell-specific vector for 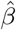.
3. We calculate the Hellinger distance between the data 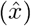 and the model predictions (*x*^**⋆**^ : *H* = *d*(*x, x*^**⋆**^)).
4. We repeat the steps above 10, 000 times, whereby we obtain a vector of distances, ***H*** = (*H*_1_, *H*_2_, …, *H*_10,000_). The parameter sets within the lowest 5 of ***s*** are accepted.
5. Lastly, we use the medians of the accepted parameter sets as point estimates for the three kinetic parameters.

#### 2.4.2 Direct likelihood-free inference based on neural networks (NN)

Recently, machine learning approaches are finding applications in likelihood-free inference [52]. In one such approach that we have recently developed [53] (here denoted by NN), we train a Bayesian neural network using sets of parameters sampled from data and their corresponding summary statistics of the simulated output. We employ a broad deep neural network with 3 hidden layers with 100 neurons each. To avoid overfitting, we invoke early stopping and dropout during the training process [54]. By also including dropout during the inference phase in tandem with the loss function presented by Gal and Ghahramani [55], approximating a Bayesian neural network, we can furthermore access the uncertainty associated with the parameter predictions. So, given summary statistics of the data, be it synthetic or real data, the trained neural network produces samples from the approximate posterior distribution.

As in the case of the rejection ABC, we simulate data by sampling gene expression counts based on the desired kinetic parameter values and subsequently downsampling the counts based on the capture efficiency. The training and test data for the NN are thus constructed analogously to the synthetic data used in the self-consistent tests in Section 3.1. For data with 500 cells, our NN is trained on 10,000 simulations drawn from the uniform Fano prior discussed in Section 2.4.3. 4000 additional samples were drawn for the validation of the network. We note that the performance of the NN is only marginally altered when reducing the training set to 1000 simulations. When dealing with data with lower cell counts, we increase the number of simulated genes in the training and validation sets proportionally to the relative decrease in the cell count. When dealing with data with, say, 500 cells as opposed to 500 cells, the number of genes in the training set is hence increased from 10000 to 100000, while 40000 rather than 400 genes are used in the validation set.

The NN draws on 15 summary statistics. Three of these measures relate to the raw reported gene expression after downsampling. These measures are the logarithm of the mean number of reported transcripts, the logarithm of the sum of reported transcripts, and the fraction of zeros in the scRNA-seq data. For the remaining twelve summary statistics, we estimate the capture efficiency based on the data and scale the reported gene expression based on this estimate. The summary statistics that rely on the scaled gene expression include the logarithm of the range of scaled transcript count, the logarithm of the highest scaled transcript count, the 10^th^ percentile, the 25^th^ percentile, the median transcript count, the 75^th^ percentile, the 95^th^ percentile, the logarithm of the variance, the skewness (3^rd^ moment), the kurtosis (4^th^th moment), the logarithm of the coefficient of variation, and the logarithm of the total number of scaled transcript counts. All logarithms are base lO. With this wide range of summary statistics, we aim to capture most of the information present in the scRNA-seq data. The compression of distributions into a set of key features thus often comes with the loss of relevant information (cf. a discussion on the sufficiency of summary statistics by [56]). However, we also note the performance is not dramatically compromised by dropping some of these summary statistics.

#### 2.4.3 Choice of priors in simulation-based methods

We need a choice of prior for the parameters of the model for the ABC method. The same priors is used to train the NN method and also to generate synthetic data for bench-marking. We have chosen these priors in a specific way to create reasonable parameter sets. In this paper, these kinetic parameters that enter Eq. (2) are computed in the following steps. First, we draw the logarithm of *k*_on_ from a uniform prior:

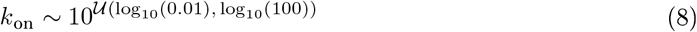

Secondly, we draw the ratio between *k*_on_ and *k*_off_ from a normal prior with a mean of 0.05 and a standard deviation of 0.5:

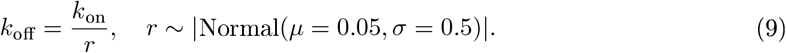

Finally, we compute *k*_syn_ based on the values of *k*_on_ and *k*_off_ as well as the Fano factor (*F* ; variance over mean, for constraining parameter sets lie in reasonable range.):

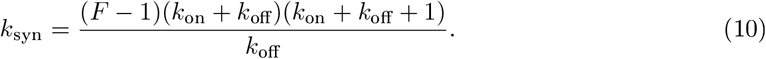

Here, we assume that the prior on the logarithm of the Fano factor is uniform: 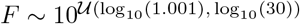.

Given a parameter set ***θ*** = (*k*_on_, *k*_off_, *k*_syn_), we simulate the outcome for a predetermined number of cells by drawing from a Beta-Poisson distribution. We then downsample the synthetic data based on the capture efficiency, *β*. When simulating synthetic data, we either employ a fixed value for all cells, mirroring the approach by Larsson et al. [42], or draw individual values for *β* for each cell from a log-normal distribution. For this purpose, we draw samples from a log-normal distribution with a mean of 2.74 and a standard deviation of 0.39 [31]. This distribution is subsequently scaled to have a mean of 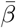.

### 2.5 Extension to non-allele-specific counts

The previous methods can be applied to allele-specific scRNA-seq data since they rely on a model for single genes based on the telegraph model. To go beyond this limitation, we modified the methods discussed above so that they can be directly applied to non-allele-specific scRNA-seq data that measures the sum of transcript counts from both alleles for each gene. To this end, we have made the simplifying assumption that the two gene alleles share the same parameters and their expression is independent of one another.

For MLE applied to non-allele-specific data, including protocols based on Unique molecular identifiers (UMI)[57], the likelihood is the distribution of observed counts of two independent integer-valued random variables (X=Y+Z), which can be obtained as follow: 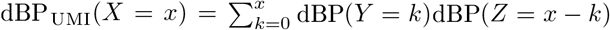. It follows that *ℓ* = Σ log(dBP_UMI_(*x*)) for optimization, where the summation is taken across cells. However, for our purposes, it is not efficient to evaluate the log-likelihood in this manner. We hence omit MLE when dealing with non-allele-specific data.

In order to apply MME to non-allele-specific counts, the procedure can be modified by replacing *M*_1_, *M*_2_ and *M*_3_ in Eq. (6) with *m*_1_, *m*_2_ and *m*_3_ defined below:

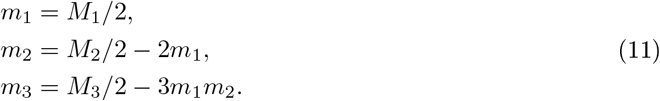

With regards to rejection ABC and the training data for the NN, we simulate non-allele-specific data by taking the sum of two random variables drawn independently from a Beta-Poisson distribution and subsequently downsampling the total count.

## 3 Results

### 3.1 Benchmarking on synthetic data

Our aim is to infer the parameters of the classic model of stochastic gene expression, the telegraph model (Fig. 1(a)), from scRNA-seq data. As illustrated in Fig. 1(b), gene expression is coupled to cell size and scRNA-seq observations are affected by heterogeneous cell-specific capture efficiencies inherent to scRNA-seq protocols. This makes inference of the parameters of the gene expression, also referred to as burst kintetic parameters in this study, from downsampled scRNA-seq data a challenging task (as illustrated in Fig. 1(c)). As discussed in Section 2, the inference methods we are considering firstly include the existing bare maximum likelihood (BMLE) and bare method of moments estimation (BMME), in which raw scRNA-seq counts are used for inference [42]. In this study, we have introduced modified MLE and MME methods (denoted simply as MLE and MME), where the cell size and capture efficiency variability are taken into account in an approximate manner (see Sections 2.3.1 and 2.3.2). We have also introduced two likelihood-free approaches, the approximate Bayesian computation rejection sampling scheme (ABC) and a direct inference approach based on the Bayesian neural networks (NN) presented by Jørgensen et al. [46] (see Sections 2.4.1 and 2.4.2).

We begin the result section by benchmarking the performance of the different inference methods on synthetic data sets that are generated from known gene-specific parameter sets as discussed in Section 2.4.3. By comparing the inferred parameter sets to the ground truth, this section thus presents a self-consistency check that allows for an evaluation of the different methods in ideal settings. The synthetic data sets include different numbers of cells, spanning from 200 to 5000. In each case, we sample between l000 and 700 different combinations of kinetic parameters, repeating each combination 20 times.

When assuming a fixed capture efficiency of l.0, we find that all methods yield accurate and precise predictions for single-allele data. We summarize the results of this analysis in Fig. S2-Fig. S4 in the supplementary material. However, this scenario is not realistic; in real-world experiments, the capture efficiency is variable and much lower than one. So, next, we created another synthetic data set for single-allele measurements with 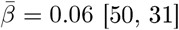. For this data set, we find that the BMLE and BMME procedures by Larsson et al. [42] lead to a pronounced systematic bias (offset) between the predictions and the ground truth for *k*_off_ and *k*_syn_ as well as the ratio of the two (Fig. 2). As a result, the scores of many performance metrics, including the mean squared and absolute errors, fall below those obtained from randomly assigning values to these parameters (Figs. S7 and S8). This makes sense as these methods effectively assume that the capture efficiency is 100% by not considering any normalisation. Moreover, both the BMLE and BMME procedures fail to attribute parameter values to a large fraction (about 65%) of the data set, yielding no parameter estimates and also producing outliers when the optimisation methods fail. We note that while the modified MLE correctly includes capture efficiencies and therefore does not suffer from the systematic bias observed in BMLE, it suffers from numerical problems in the evaluation of the modified likelihood and the optimisation (see Figs. S5 and S6). In contrast, our simulation-based approaches, rejection ABC and the NN, consistently yield accurate and precise predictions across all data sets and kinetic parameters. They thus consistently yield the lowest mean absolute and mean squared errors among all six methods, and the true values lie within the assigned 95% confidence interval of both methods in the majority of cases (Fig. 2).

**Figure 2:**
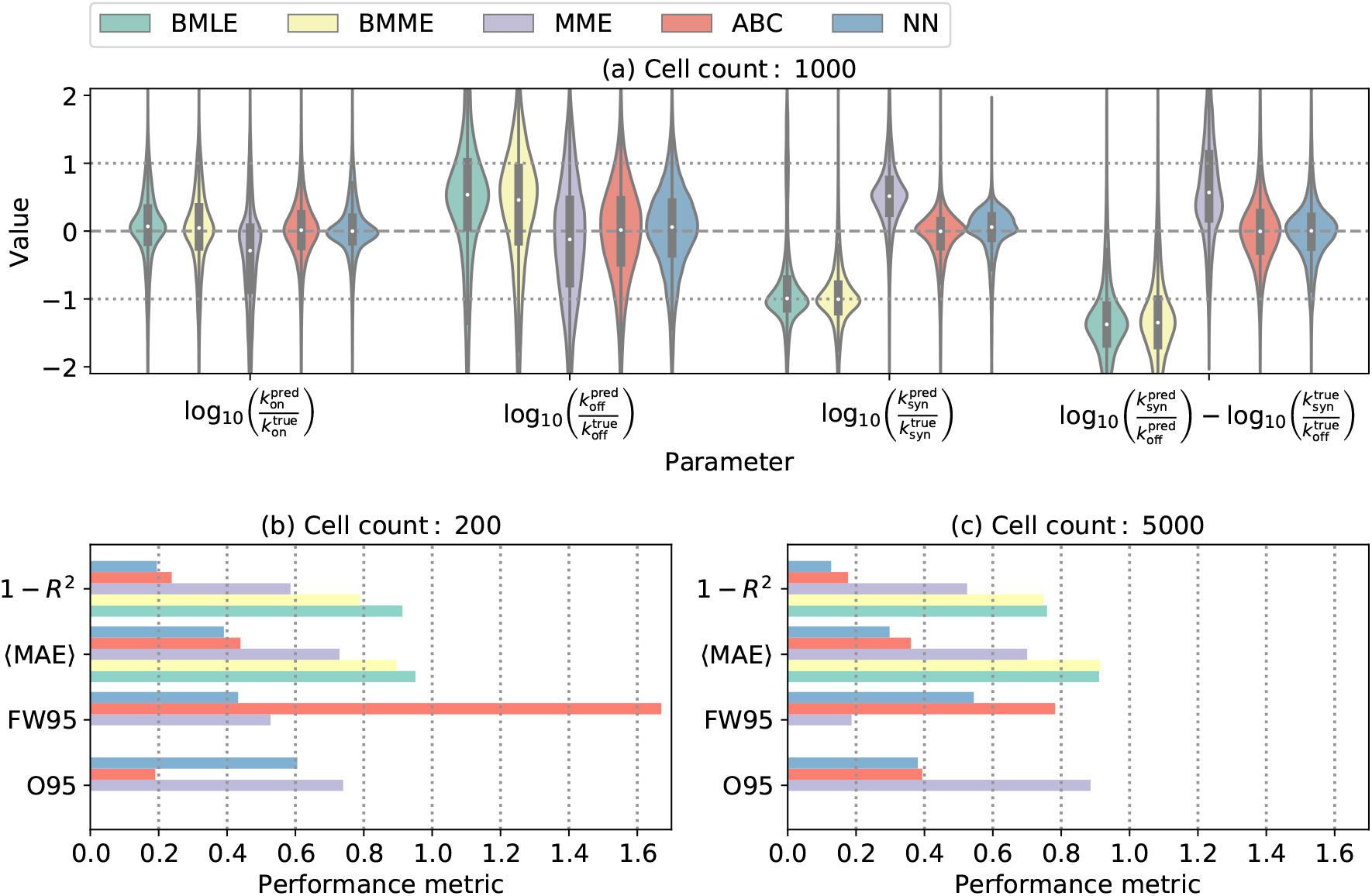
Comparison between different modelling approaches allele-specific synthetic data with 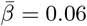. **(a)**: logarithmic residuals across all four parameters for a data set containing 1000 cells. **(b-c)**: contain four performance metrics for data containing 200 and 5000 cells, respectively. These metrics are the coefficient of determination (*R*^2^), the mean absolute error (MAE), the fraction of the true parameter values that lie outside of the 95 % confidence intervals (095), and the width of the 95 % confidence intervals in logarithmic space (FW95). 0nly the NN, ABC and MME supply confidence intervals. For each number of cells, the synthetic data set contains 7000 genes with 20 repetitions each. All metrics (except for FW95) are formulated such that a lower value implies a better fit. Note that the modified MLE is omitted from this summary as our implementation suffers from numerical issues (see Fig. S5).

As seen in Fig. Fig. 2, the accuracy of inferring *k*_*off*_ is the poorest among the kinetic parameters, suggesting some degree of non-identifiablity. Also, as expected, increasing the number of cells from 2OO to 5OOO improves the performance metrics of all methods. Interestingly, NN has the best performance at small cell numbers. We also note that only the MME, ABC and NN attribute confidence intervals while the remaining methods solely provide the best fit (Figs. S4 and S8). The MME generally leads to narrower confidence intervals than both the ABC and NN, but a significantly larger fraction of the true values do not lie within the error bars of the MME, suggesting that the MME significantly underestimates the prediction error.

Finally, we developed a modified MME, ABC and NN method that works for non-allele-specific data (Section 2.5) and benchmarked their performance on synthetic non-allele-specific data. We find that the NN yields narrower and smaller residuals. The results are summarized in Fig. S9 in the supplementary material. So, overall, we propose that the NN method is the most robust approach, and we mostly use this approach in the applications to real data in the rest of this study.

### 3.2 Sparsity of gene counts lead to wrong model identification

We note that even if expression counts are drawn from Beta-Poisson distribution, they may equally well fit by other distribution models depending on parameters. This can be the case, for example, if these models have fewer parameters such as the Poisson distribution or the negative binomial (NB) distribution which have one and two parameters, respectively, compared to the three parameters of the Beta-Poisson model. To investigate this we use Akaike information criterion (AIC), which is a commonly used metric for model selection that accounts for both quality of the fit (likelihood of data) and the complexity of the model (number of parameters). We generated a simulated data set (5OO cells and 7OOO genes) using the Beta-Poisson model and we calculated the AIC using the following three models and choice of parameters for each gene:

- Beta-Poisson: ground truth parameters were used.
- Negative binomial: R package bayNorm[31] (NB model for non-allele specific scRNA-seq data) was applied to the raw counts to infer NB parameters for calculating the AIC.
- Poisson: raw counts were scaled by 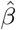, then the mean expression of each gene was calculated. For each gene in each cell, the mean expression was multiplied by 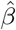 to be the mean parameter in the Poisson distribution, which was used for calculating the Poisson model AIC.

Then genes were assigned to one of three models (Beta-Poisson, negative binomial or Poisson) with the lowest AIC value. As data is generated from a Beta-Poisson model, one might expect this model always selected, however, we found for many genes one of the simpler models are selected. The genes in the Poisson and NB category tend to have the lower mean expression (Fig. 3(a)), which highlights the fact that there is less information for estimating Beta-Poisson parameters. Indeed, *k*_syn_, which regulates the mean expression, has the highest impact on the identifiability of the Beta-Poisson model Fig. SlO. This indicates that inference of burst kinetics is only possible for genes that have high enough expression as expected. In line with this result, we observe that the inference accuracy is poorer for the lowly expressed genes in our synthetic data (Fig. 3(b), Fig. S11).

**Figure 3:**
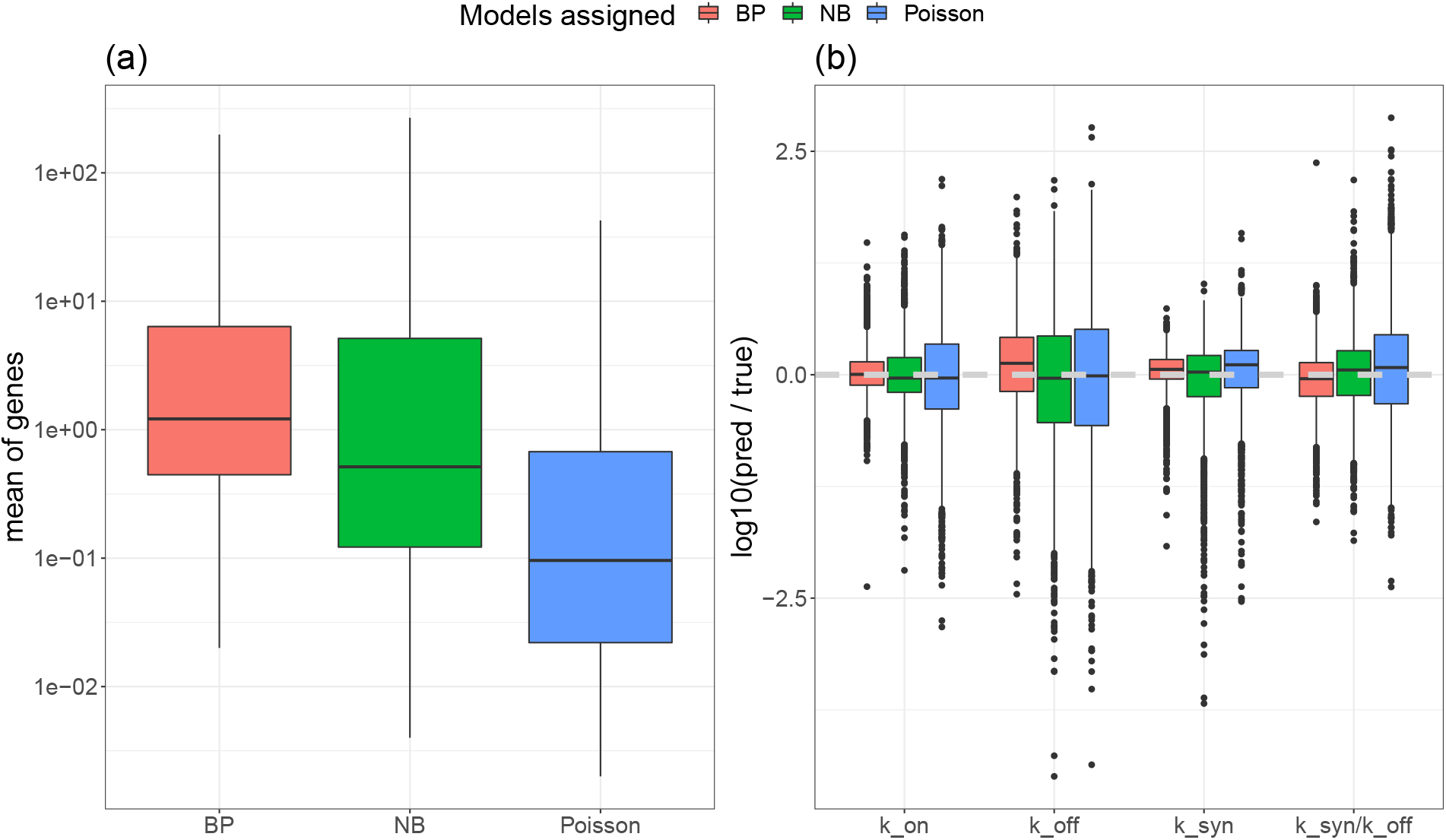
Genes with low counts are assigned to simpler models. **(a)** Based on synthetic data generated by the Beta-Poisson model, genes were labelled to be from one of the three models according to the their AIC value. The mean counts for genes assigned to each model is shown. **(b)** The ratios between inferred and true parameter values in each group of genes are shown. Estimates from genes which are assigned to BP correctly are closer to ground truth values.

### 3.3 Application to real-world data

#### 3.3.1 Estimating kinetic parameters from individual allele data

We used the NN method to reassess the allele-specific data from Larsson et al. [42] containing lO727 genes and 224 cells. The data contains missing values. The number of missing values varies between different genes. Here, we only include genes with mean expression across non-missing values above 1. As shown, this is important as genes with low counts do not contain enough information. This first filtering leaves us with 1992 genes. Of these genes, we remove genes with large number of missing values. This leaves us with 1953 genes. We find that the NN yields kinetic parameter estimates that are consistent with those obtained from the BMLE procedure by l29j when assuming that 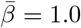. However, as seen in Fig. S12 using realistically small and cell specific capture efficiencies leads to a systematic shift to higher burst sizes and a wider spread in burst frequency. The choice of prior used in training our NN inference method has only a small effect on the inference results, which suggests the robustness of our method Fig. S12.

As investigated in the original study [42], we look at the link between the presense of TATA elements and Initiator (Inr) and the burst kinetics using our inferred parameters. We find NN yields kinetic parameter estimates that are qualitatively consistent with those obtained from the original MLE procedure by [42] such that genes with TATA elements have larger burst size (Fig. 4 (a-b)). By filtering out lowly expressed genes, our analysis reveals that genes with only Inr can boost burst sizes (Fig. 4). Similar qualitative results can be achieved via the MLE approach adapted by Larsson et al. [42] after removing the lowly expressed genes (Fig. 4). In addition, the NN results reveal that genes with Inr have lower burst frequency than genes without TATA element and Inr. We note that Larsson et al. [42] did not arrive at the same conclusions. The explanation for this is that they kept around 7000 genes which include lots of lowly expressed genes for which inference results are not reliable (see Section 3.2 and Fig. S11). Interestingly, the NN results also reveal that genes with both TATA and Initiator (Inr) tend to have relatively lower burst frequency compared with genes without TATA and Inr, which was not found by [42] (Fig. 4 (d)).

**Figure 4:**
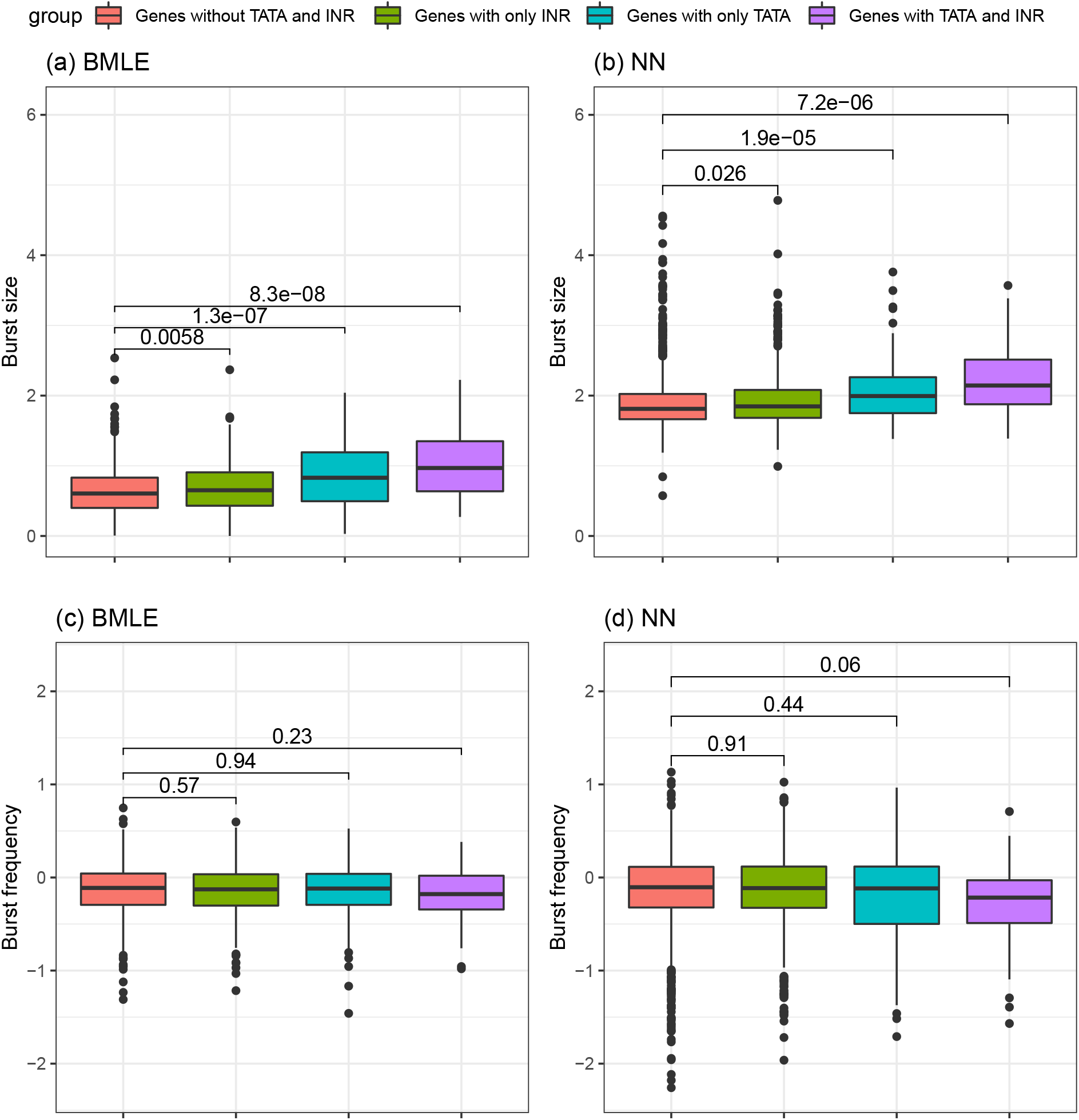
The relationship between burst kinetics and promotor characteristics based on the NN results with 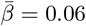 and the BMLE from Larsson et al. [42]. Results are shown for Allele c57. **(a-b)** Box plots of Burst size and **(c-d)** burst frequency estimates of genes with or without TATA elements and Inr. The P-values of the Wilcox test between groups are shown.

Based on the present data set, we note that we find the simulations to successfully recover the observed relation between the dropout rate and the mean expression for each allele (Fig. S14), providing further support for the accuracy of our mathematical model of scRNA-seq data.

Finally, in our inference methods, the only source of biological and technical variability is considered to be the cell size and capture efficiency. To test the validity of this assumption in real data, we produced synthetic data for two independent alleles from 100 cells down-sampled by the same capture efficiency using parameters inferred by the NN method on the single-allele data of Larsson et al. [42]. We then computed the correlations between the two alleles in our synthetic data and plotted the results against the correlation between two alleles in the real data (Fig. S15). We find a clear linear relationship between the simulated correlation and real correlation. This indicates that it is reasonable to consider cell size and capture efficiency as an important source of extrinsic noise since we observe most genes to have a significantly positive correlation between the two alleles, captured in our simulations. These results suggest the observed positive correlation between the gene expression between the two alleles can be well explained by variation in capture efficiency across cells. So, one does not need to invoke correlated activity between the alleles or other significant sources of extrinsic noise. This further motivates the approach we have proposed for the inference of gene expression parameters from non-allele-specific data. Interestingly, for some genes in the real data, there is a negative correlation between two alleles, which might indicate anti-correlation in the activity of those genes.

#### 3.3.2 Estimating kinetic parameters from non-allele-specific scRNA-seq data

In this section, we analyse scRNA-seq data of mouse brain cells form two recent study (Mizrak et al. [58] and Ximerakis et al. [59]) to highlight the application of our inference methods (MME, ABC and NN) on non-allele-specific data that assumes that the counts are related to the sum of two identical but independent alleles (Section 2.5).

The data from Mizrak et al. [58] contains 28407 cells from mouse brains (after removing doublets), and covers multiple cell types like neuronal progenitors (active neural stem cells, transit amplifying cells, and neuralblasts (aNSC+TAC+NB)), oligodendrocyte progenitor cells (OPCs), committed oligodendrocyte precursors (COPs), oligodendrocytes (OLG), microglia (MG), astrocytes (ASC) and neurons. In addition, we explored the data from mouse brains Ximerakis et al. [59], where there are 37069 cells collected from either young or old mice. The data set contains various cell types, including Neural stem cells (NSC), mature neurons (mNEUR), OPC and other cell types from young and old mice.

Cell type markers are by definition the ones that are overexpressed in a particular cell type but not others. Here, we investigate if these gene expression alterations are associated with the changes in bust size or burst frequency. When comparing stem cells (aNSC+TAC+NB) with other differentiated cells, all inference methods reveal higher burst frequencies for stem cell markers in stem cells than differentiated cells like neurons and oligodendrocytes (Fig. 5). To a lower degree we see burst size (Fig. S16), though burst size from MME is not consistent with ABC and NN. Interestingly, cells at different stages of oligodendrocyte differentiation (COP, OPC and oligodendrocyte) tend to have either slighterly higher or similar burst frequency/size to the other mature cell types.

**Figure 5:**
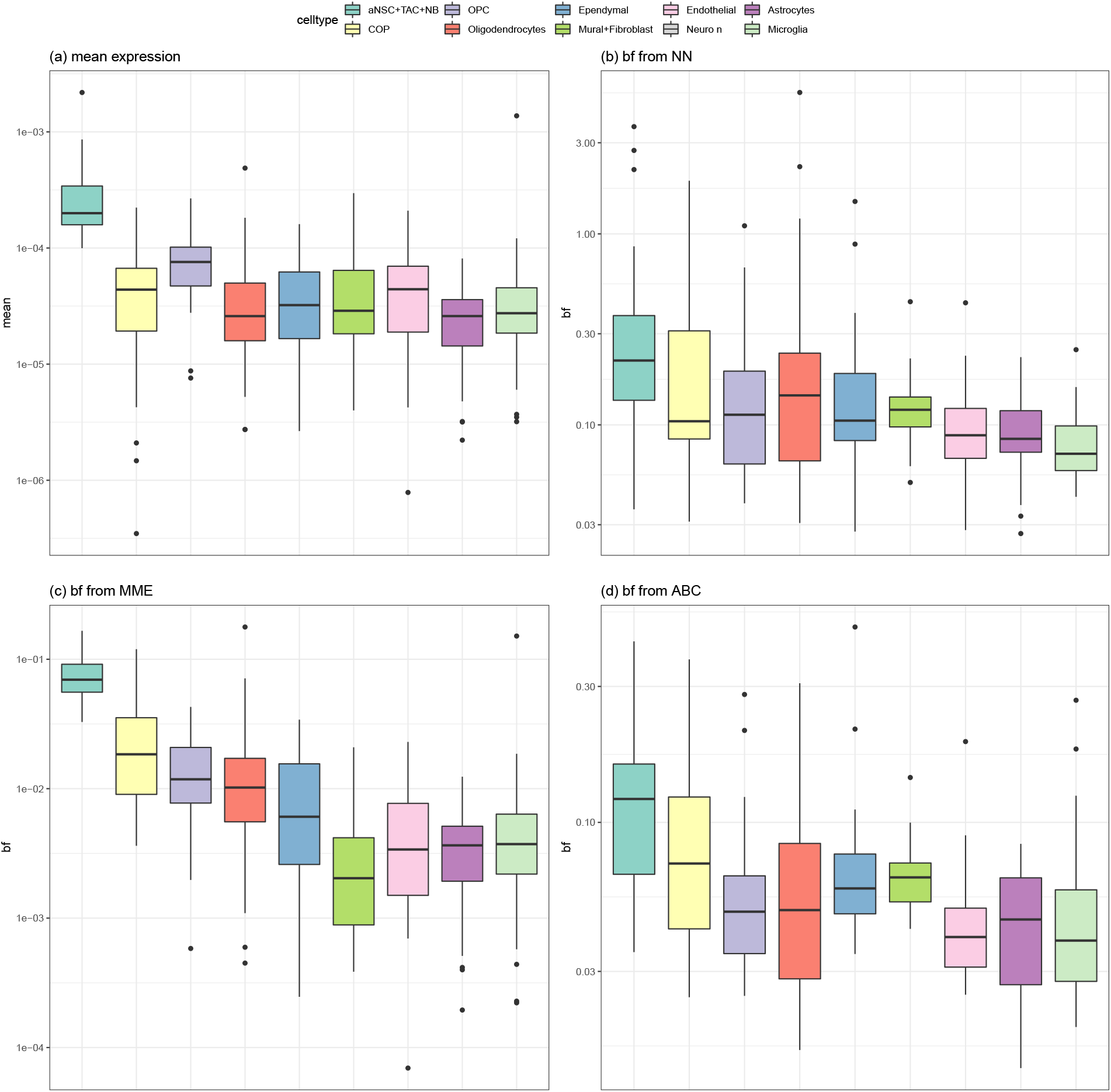
Higher expression of stem cell marker genes in stem cells are associated with higher burst frequency. Cell types are allocated using aNSC marker genes reported in Mizrak et al. [58]. The first three cell types are stem cell-like and the rest are mature cell types **(a)** Box plots of mean expression of stem cell markers across the cell types are shown. Mean expressions was calculated after total count normalized; Box plots of inferred burst frequencies using NN **(b)**, MME **(c)** and ABC **(d)** inference approach.

Our second dataset from Ximerakis et al. [59] that has data from both young and old brains confirms the high burst frequency for stem cell markers in stem cells regardless of brain age (Fig. S17). In this study, they have reported that genes encoding ribosomal subunits have a reduced expression upon aging [59]. We asked is it burst frequency or burst size of the ribosomal genes that is down regulated upon aging. Results from NN and ABC shows again burst frequency but not burst size is downregulated for ribosomal genes upon aging in NSC[59], ASC[59, 60] and OPC[59] compared with other mature cells (Fig. 6).

**Figure 6:**
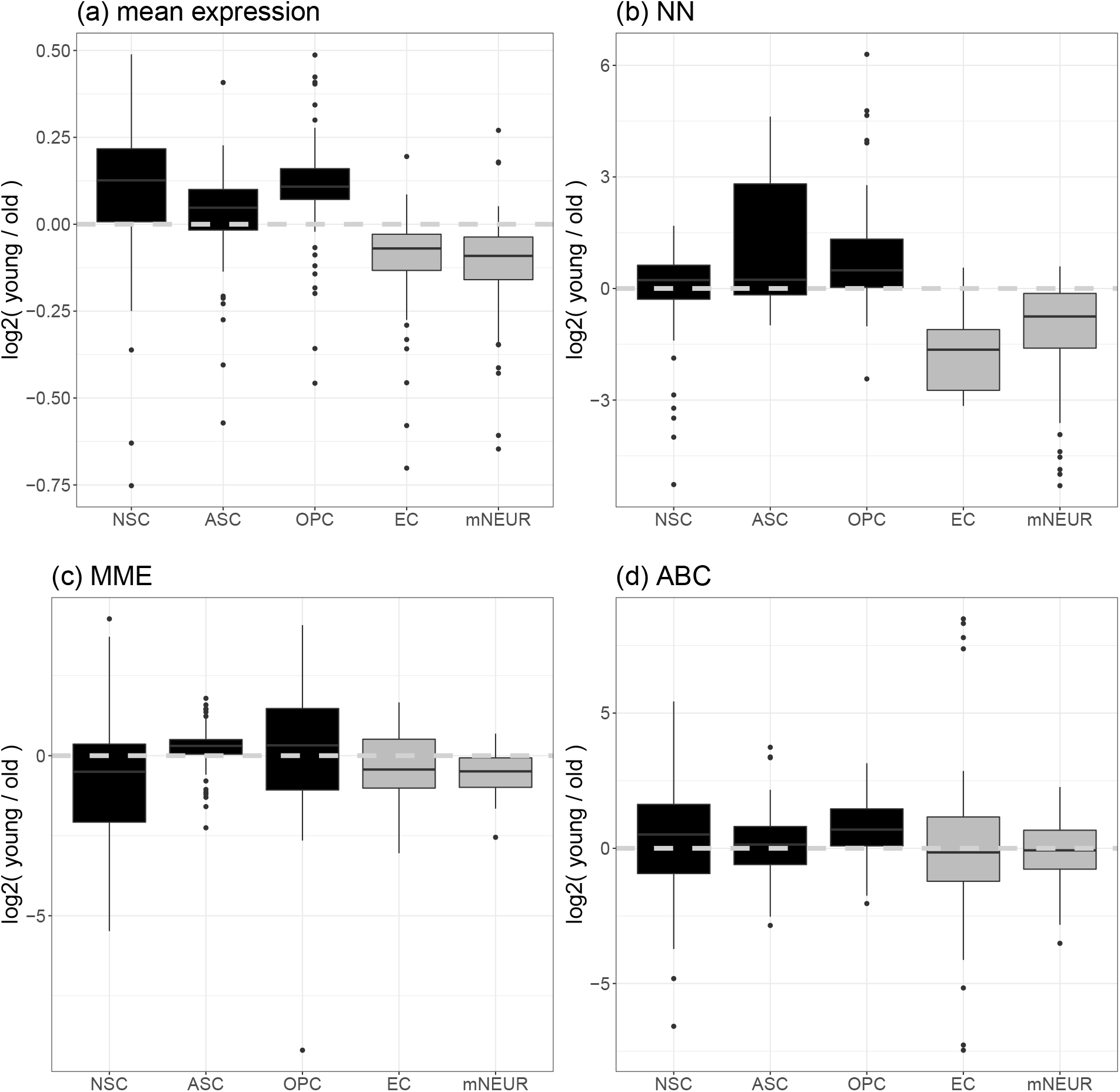
Ratio of burst frequency of genes encoding ribosomal subunits from young and old mice within each cell type. Dark color shows the ratios within NSC and progenitor cells; light color shows the ratios within mature cells. **(a)** Box plots of ratio of mean expression of genes encoding ribosomal genes. Mean expression was calculated after total count normalized; Box plots of ratio of inferred burst frequencies using NN **(b)**, MME **(c)** and ABC **(d)** inference approach.

## 4 Discussion

In this paper, we revisited the problem of inferring burst kinetics of gene expression from scRNA-seq data. We provide a novel expression for the likelihood for single-allele scRNA-seq data, which allows us to take cell-to-cell variation in size and capture efficiency correctly into account. We show that maximum likelihood estimation (MLE) could be unreliable due to numerical challenges. However, we introduce likelihood-free approaches that include a modified method of moments (MME) and two simulation-based inference methods. Through a series of benchmarks on synthetic and real data, we demonstrate the reliability and flexibility of the simulation-based inference methods. We show that these methods also provide confidence intervals and could also be easily generalised to non-single-allele situations, which makes them more widely applicable. We obtain the best results using a recent implementation of a simulation-based inference method based on Bayesian neural networks [46].

Recent studies have used the maximum likelihood estimation method using a Beta-Poisson model without any normalisation Larsson et al. [42], Kim and Marioni [25] termed here as bare maximum likelihood estimation (BMLE). As we show in this paper, this approach can result in biased and distorted distributions of estimates for burst kinetic parameters, including burst size. Also, we show that burst kinetics parameters become unidentifiable for lowly expressed genes and that this property could result in misleading results. While maximum likelihood estimation has good theoretical guarantees, computational challenges in evaluating the likelihood and also challenges in optimisation can make this method less favourable. Indeed, a recent study has likewise highlighted the challenges with maximum likelihood estimation and the non-identifiability for similar models of stochastic gene expression [33].

There are not many available allele-specific scRNA-seq data, but UMI-based non-allele-specific scRNA-seq data are highly abundant. We have therefore modified the MME method and also our simulation-based methods to infer the kinetic parameters directly from non-allele-specific (e.g. UMI) count matrices. Although we assume that the two gene copies have identical kinetic parameters and transcribe independently in this study, we note that these assumptions can easily be relaxed for simulation-based methods. Indeed, some recent studies have suggested there is evidence for allelic imbalance and dependence in burst kinetics across the gene alleles in existing the scRNA-seq data Choi et al. [61], Mu et al. [62]. We applied our methods to two mouse brain scRNA-seq datasets. Our results indicate that gene regulation across stem cells and aging brain tends to be associated with the regulation of burst frequency and to a lower degree burst size. A recent study has proposed epigenetic regulation of burst frequency in fitness genes upon stress could underly evolution of cancer [63].

We note here we are neglecting other possible sources of extrinsic variabilities, such as fluctuations in the kinetic rates due to fluctuations of other molecules in the cells. However, we have shown here that a lot of gene expression correlations between alleles can be explained by accounting in variation in cell size and capture efficiency. In fission yeast, we have shown previously accounting for cell size variation can capture most of the extrinsic variability observed in gene expression [17]. Other studies have included the effect of different cell cycle stages, replication and gene copy numbers [33]. Sun and Zhang [57] used allele-specific expressions in diploid cells and intrinsic and extrinsic noise decomposition to study the genetic factors affecting gene expression noise. We also note that more detailed mechanistic models of RNA-sequencing protocols can help to explain more of the technical noise and biases in the data [14, 64, 65, 66, 67].

Inferring kinetic parameters of stochastic gene expression from scRNA-seq data is challenging. First and foremost, the data is sparse and has missing values. This characteristic of the data presents an obstacle to any attempt to accurately estimate the parameters. In addition, the extrinsic variables, such as cell size and capture efficiency, are usually not known (for an exception, where cell size has been measured along with scRNA-seq see [49]). Furthermore, measurements or theoretical considerations that constrain the range within which the kinetic parameters lie are not readily available. Statistical analysis, such as the one presented in this paper, would thus benefit from additional measurements or other constraints that would provide tighter priors. While many researchers have already studied the inference of kinetic parameters from high-throughput data, such as scRNA-seq data, several aspects are hence, by far, not fully explored. An important area of future research is using multi-omic single-cell data. The data is quickly becoming available and could thus inform our understanding of global gene expression variability [68, 69]. Some research is already starting in this important area based on both statistical data integration [68, 70, 71] and model-based inference [72, 73, 66]. Ultimately, by harnessing gene-gene correlations such multi-omic single cell datasets could be used for the inference of genetic networks [74, 75].

In summary, we have proposed a simple and accurate method to take into account the variation of cell size and capture efficiency in the inference of burst kinetics from scRNA-seq data. We provide likelihood free implementations of our approach that is robust and flexible and apply it to synthetic and real data. Our analysis shows how state-of-the-art inference tools can help us to extract the valuable information that is not caught by the standard approaches.

## Code availability

Code for running ABC and NN methods can be found in https://github.com/WT2l5/Julia_ABC and https://github.com/WT2l5/nnRNA_respectively.

## Acknowledgement

We acknowledge Ioannis Loukas and Paola Scaffidi for early discussion on challenges of inference of burst kinetics from scRNA-seq data. We acknowledge Dimitris Volteras for providing detailed comments on the manuscript. This work was supported by The Oli Hilsdon Foundation through The Brain Tumour Charity, grant number (GN-000595) in connection with the program “Mapping the spatio-temporal heterogeneity of glioblastoma invasion”; by a UKRI Future Leaders Fellowship (grant no. MR/T0l8429/l) to PT and by a Engineering and Physical Sciences Research Council (EP/NOl4529/l) to VS.

## A Supplementary figures

**Figure S1:**
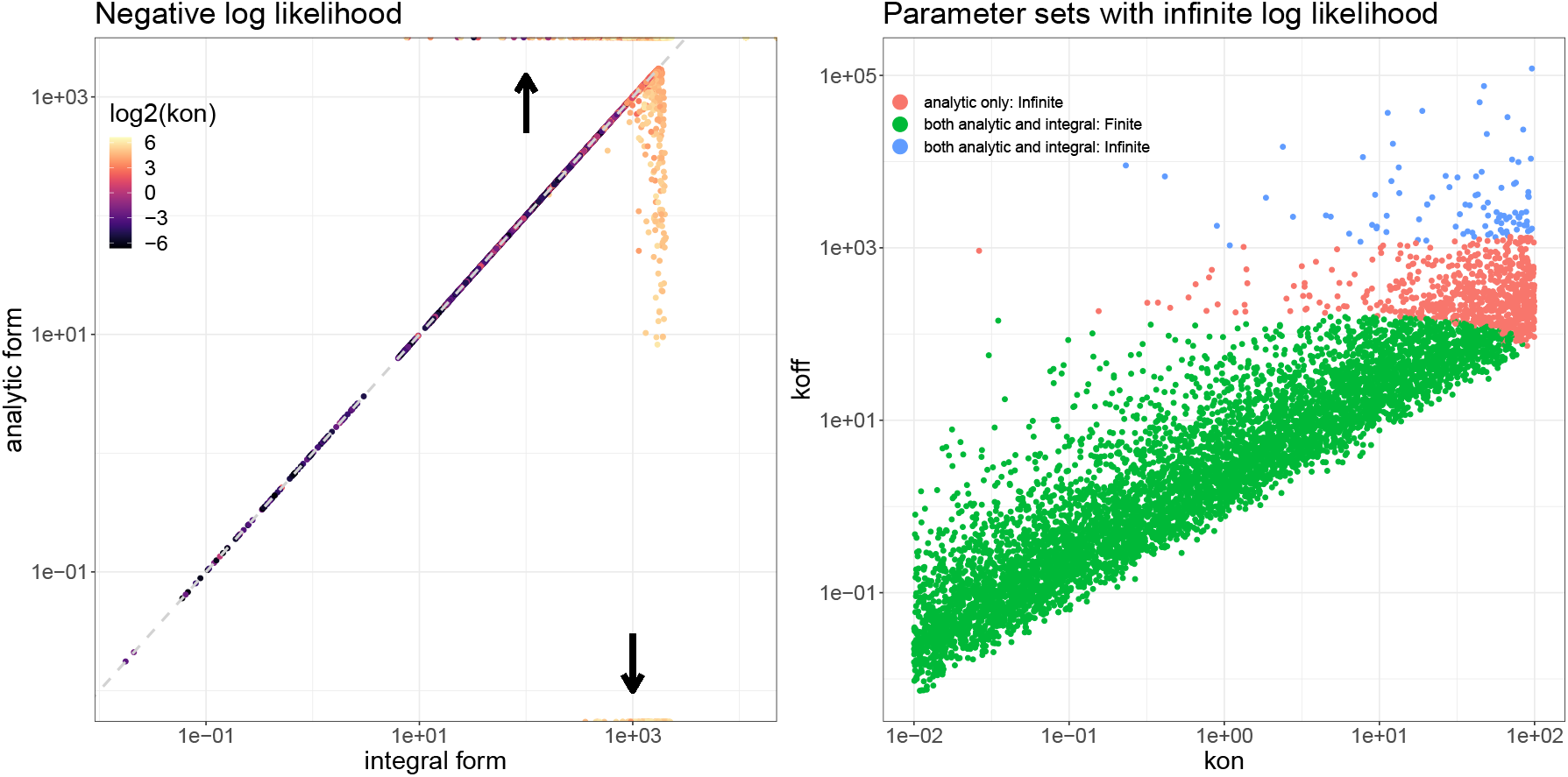
Two ways of calculating the negative log likelihood. Using the simulated data and the corresponding grosund truth parameters, the negative log-likelihoods of most genes are almost the same whether compute them using the integral method or the analytic form given in (Eq. (3)). However, the analytic form is more likely to give an infinite outcome as the upper bound and 0 as the lower bound than the integral method. The arrows in the first panel highlight this property. The inconsistencies between the two approaches stem from parameter sets where the *k*_on_ is high.

**Figure S2:**
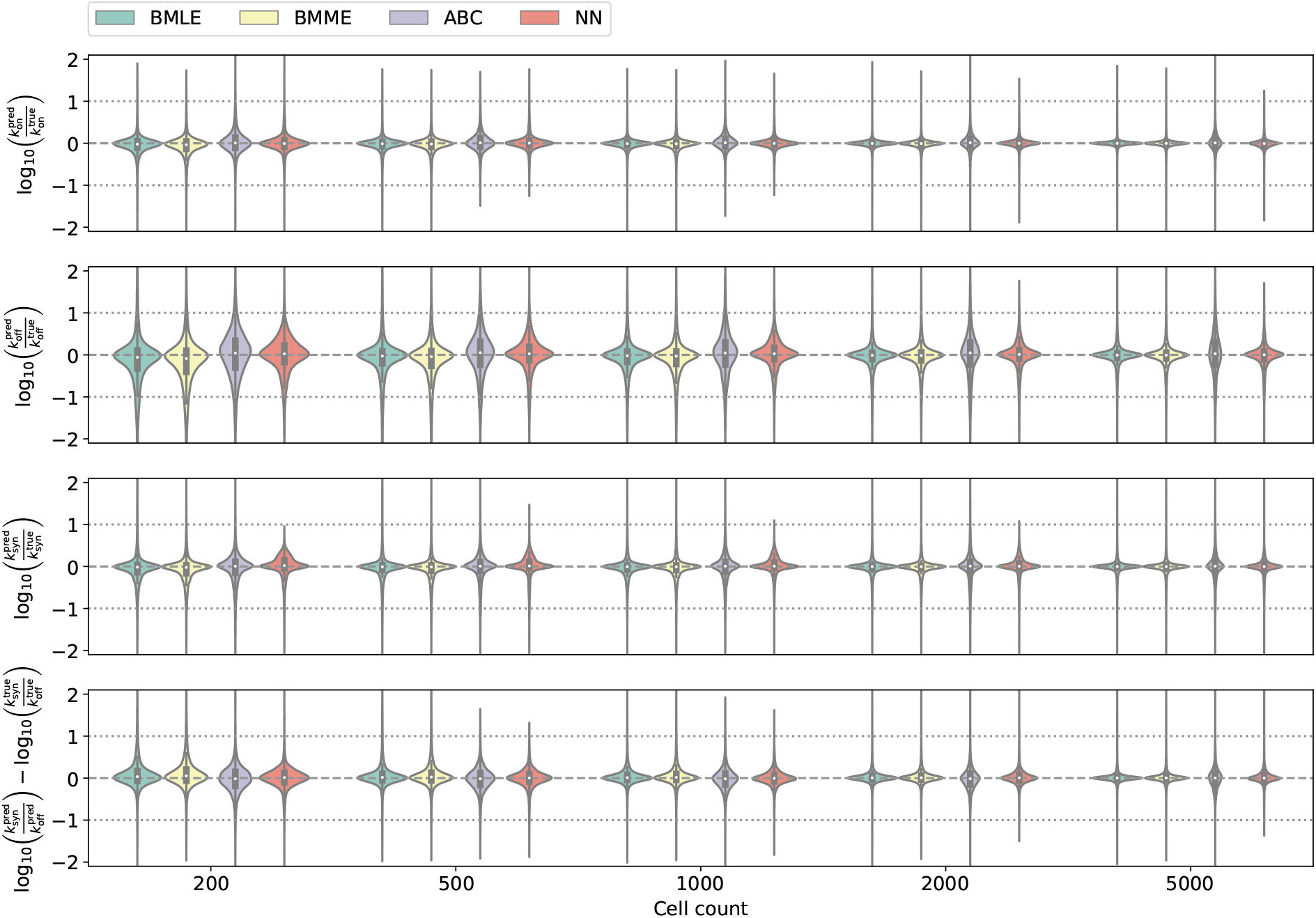
Comparison between different modelling approaches as a function of the number of cells at a fixed capture efficiency of 1.0 for allele-specific synthetic data. For each number of cells, the synthetic data set contains 1000 genes with 20 repetitions each.

**Figure S3:**
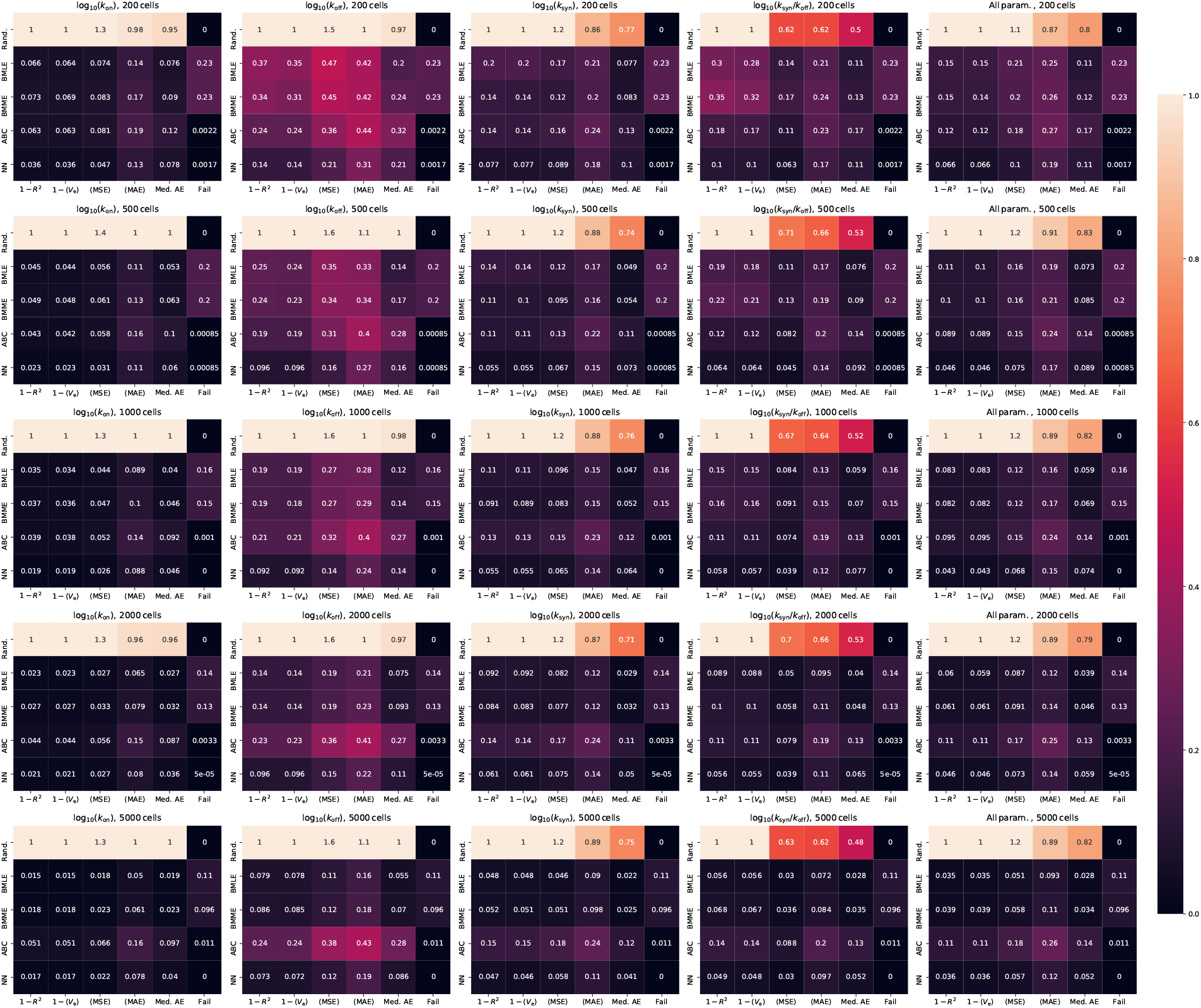
Performance metrics across methods and cell counts based on allele-specific synthetic data with a fixed capture efficiency of 1.0. For all metrics lower numbers imply better performance. These metrics include the coefficient of determination (*R*^2^), the explained variance (*V*_e_), the mean squared error (MSE), the mean absolute error (MAE), the median absolute error (med. AE), and the failure rate, i.e. the fraction of test cases, for which the method is unable to provide parameter estimates. To put the scores into perspective, the upper row of each heat map includes the results (rand.) that are obtained when consistently guessing the parameters to take the mean value of the ground truth across all samples. For all approaches, we use a uniform prior for the logarithm of the Fano factor.

**Figure S4:**
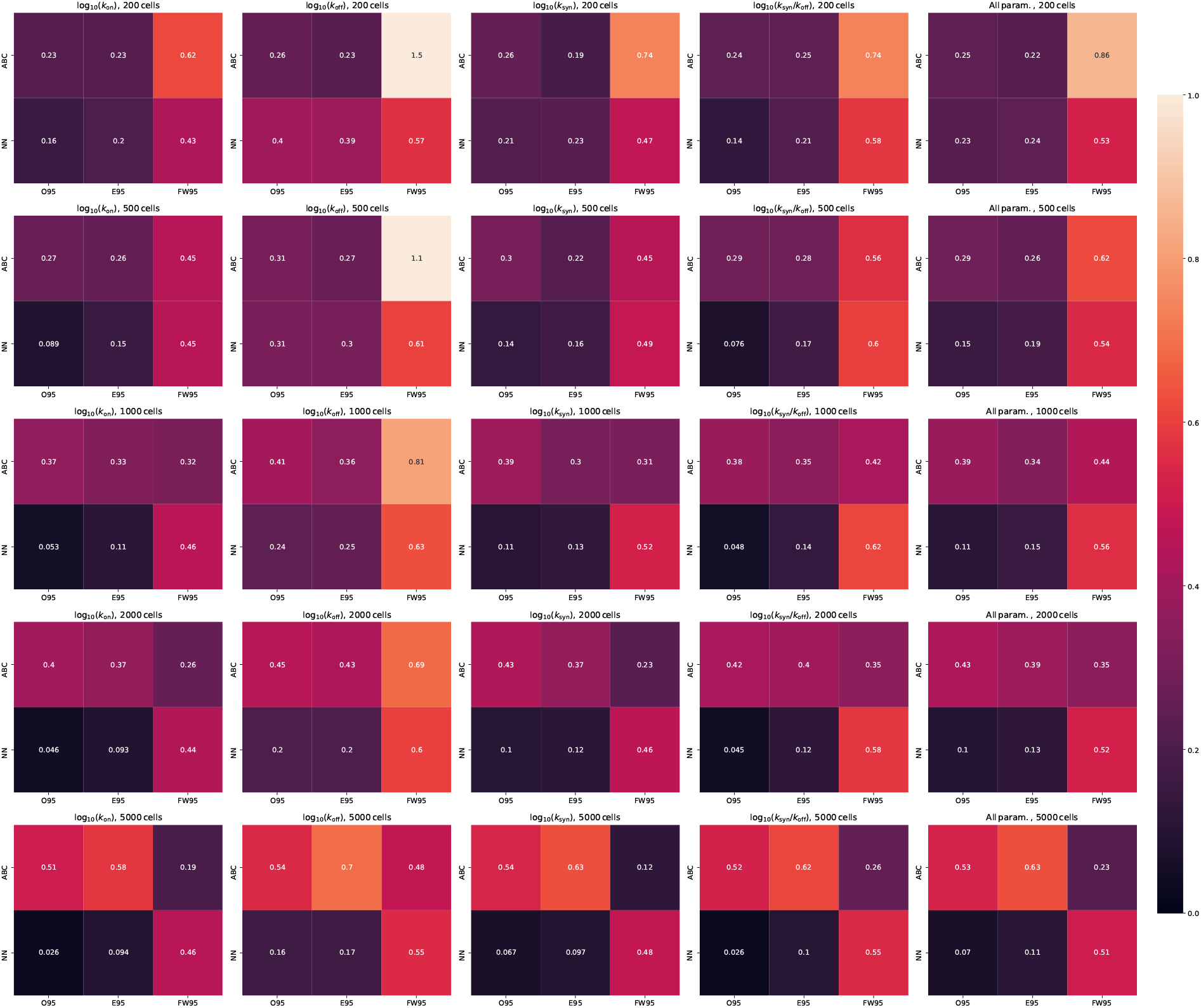
Additional performance metrics across methods and cell counts related to the predicted 95 % credibility intervals. The plot is based on allele-specific synthetic data with a fixed capture efficiency of 1.0. 095 denotes the fraction of the true parameter values that lie outside of the 95 % credibility intervals. FW95 denotes the width of the 95 % credibility intervals in logarithmic space, while E95 is the median absolute error of the predictions in units of FW95. A uniform prior for the logarithm of the Fano factor was employed.

**Figure S5:**
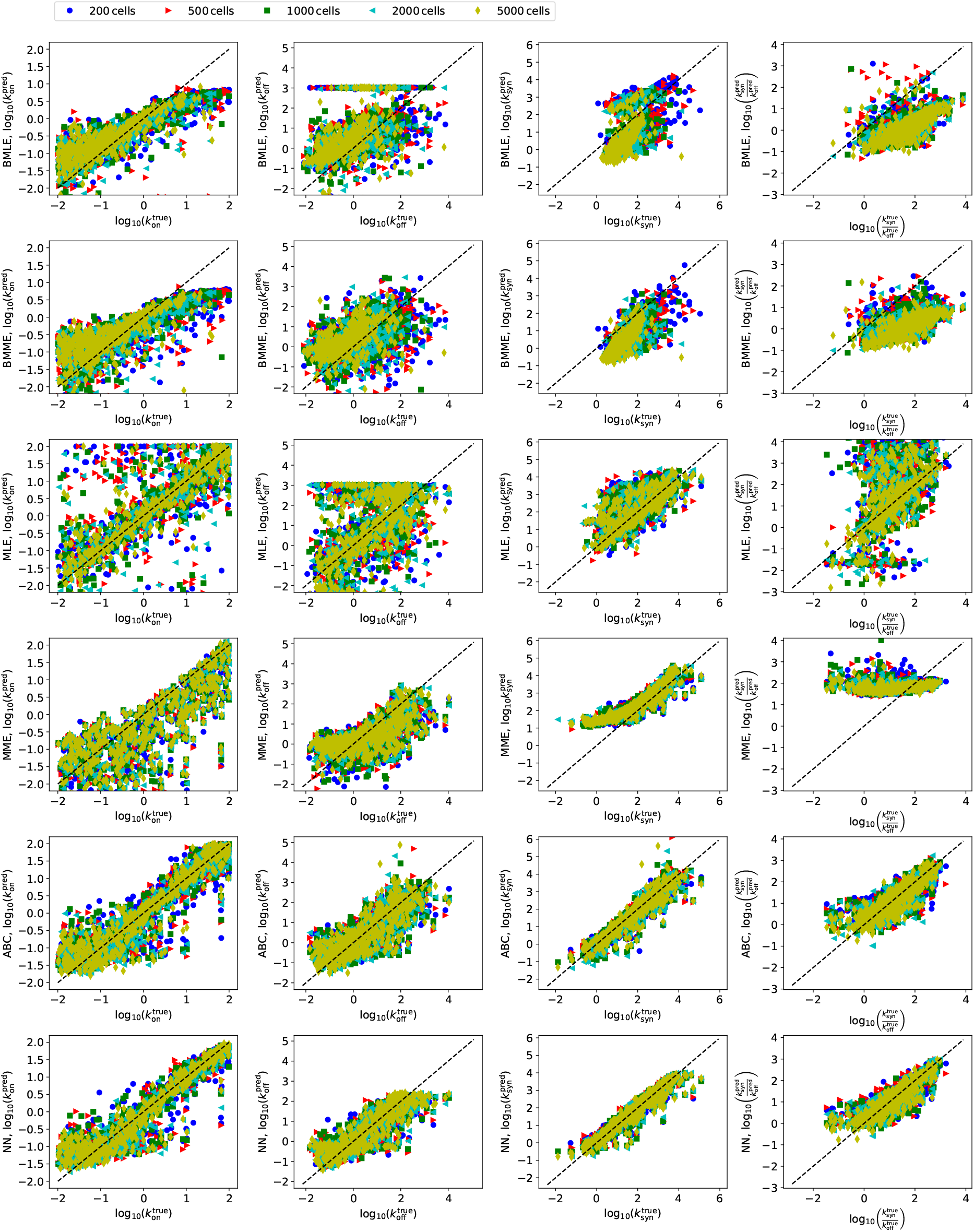
Scatter plot showing the predicted parameters as a function of the ground truth across different methods and cell counts. The plot includes 500 cells for each method and is based on allele-specific synthetic data.

**Figure S6:**
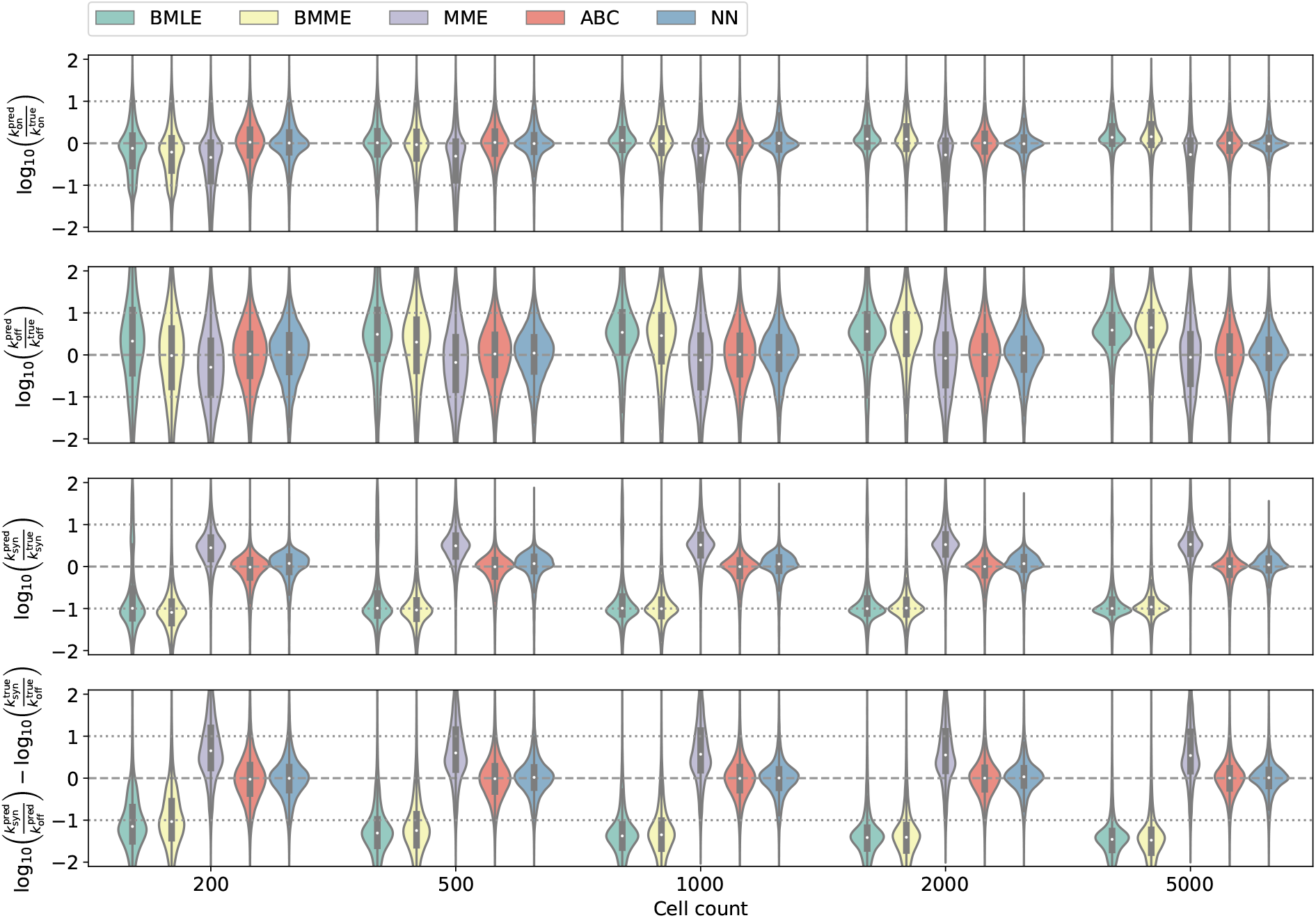
Comparison between different modelling approaches as a function of the number of cells for a varying capture efficiency with 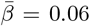 for allele-specific synthetic data. For each number of cells, the synthetic data set contains 7000 genes with 20 repetitions each.

**Figure S7:**
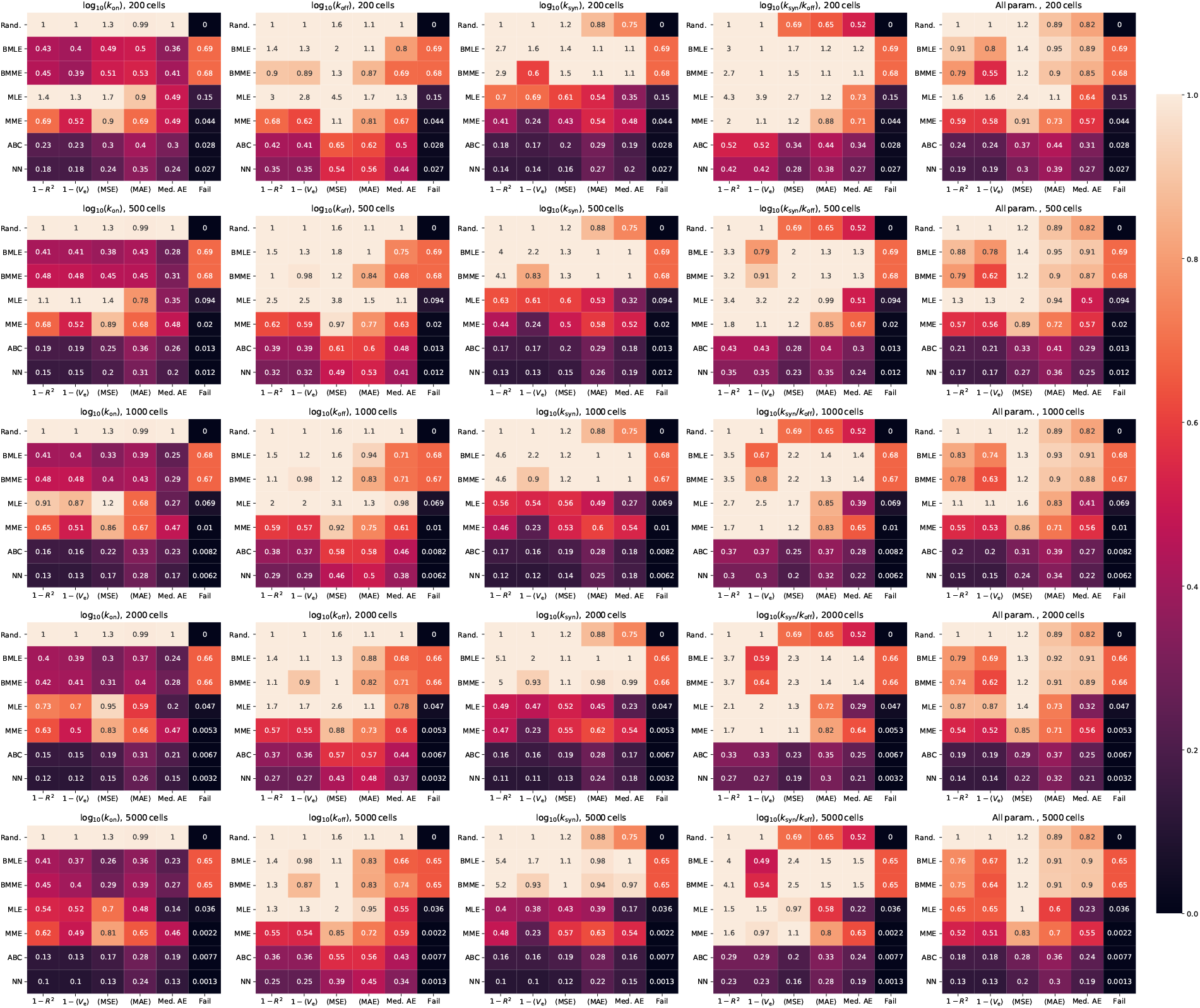
Performance metrics across methods and cell counts based on allele-specific synthetic data with a varying capture efficiency 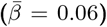. For more information, see the caption of Fig. S3.

**Figure S8:**
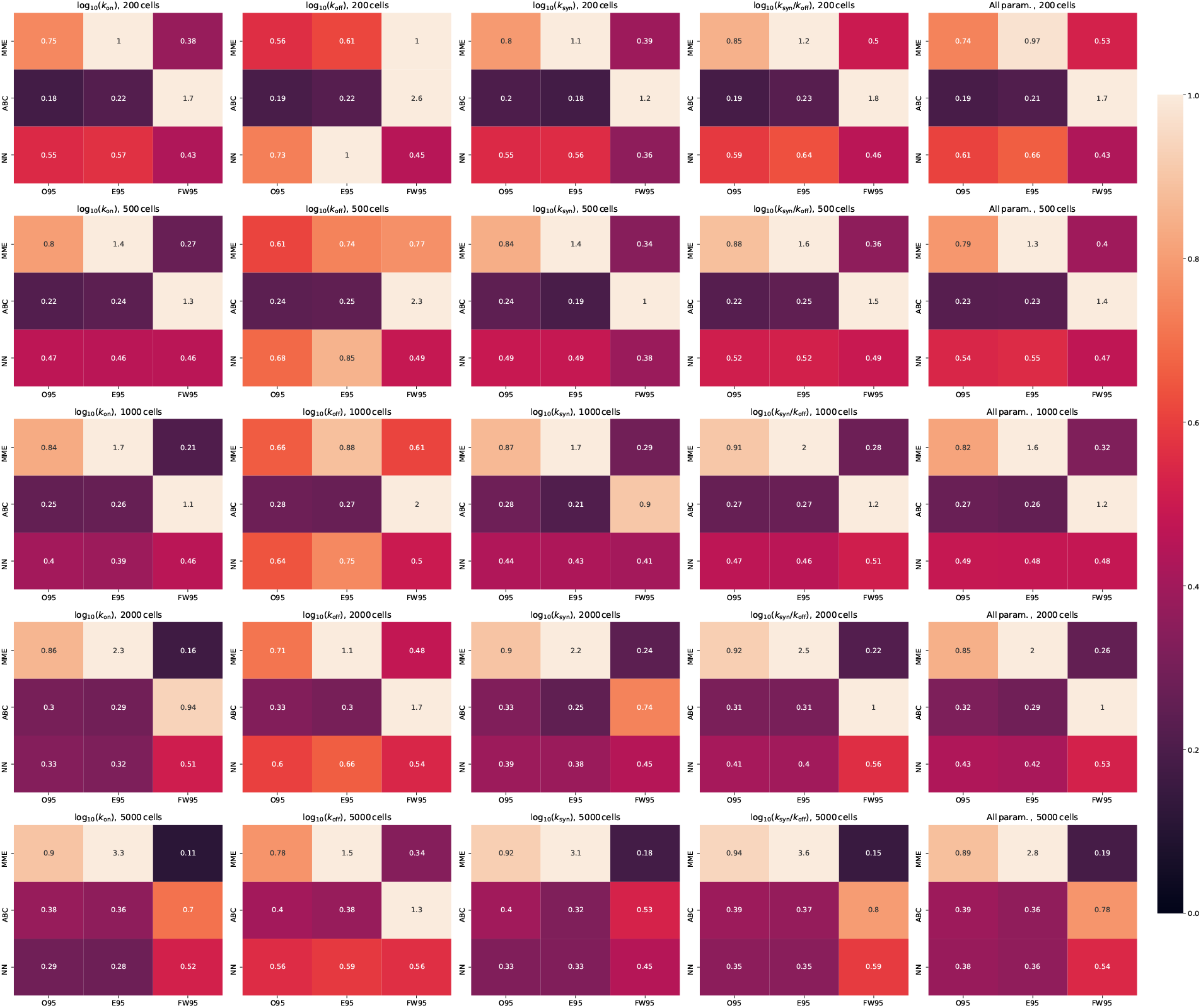
Additional performance metrics across methods and cell counts related to the predicted 95 % credibility intervals. The plot is based on allele-specific synthetic data with a varying capture efficiency 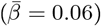. For further details see Fig. S4.

**Figure S9:**
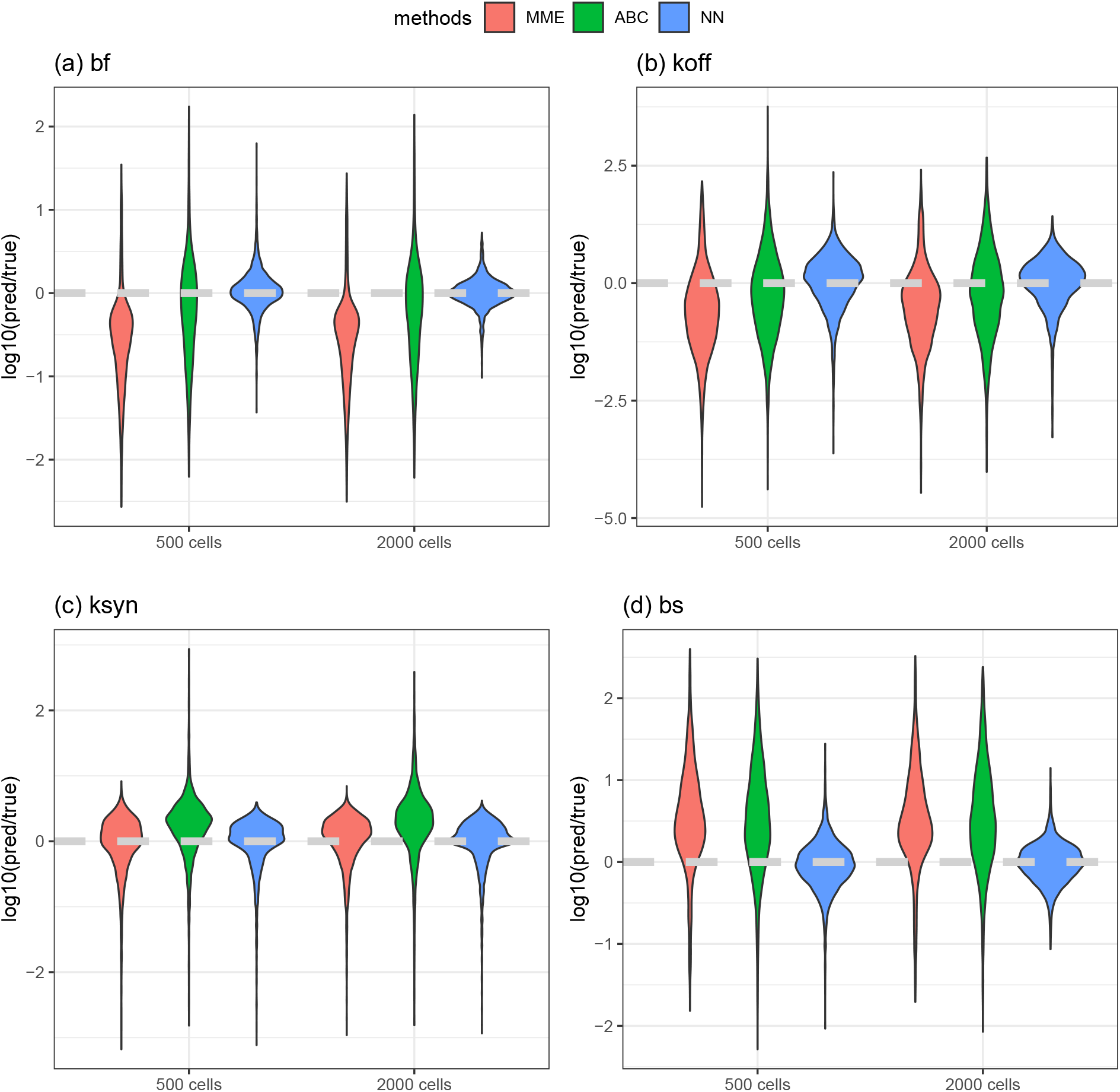
Comparison of inference methods. Based on non-allele specific simulated data with mean capture efficiency set to be 0.06. Ratio between estimates and ground truth of burst frequency **(a);** koff **(b)** ; ksyn **(c)** and burst size **(d)** are shown.

**Figure S10:**
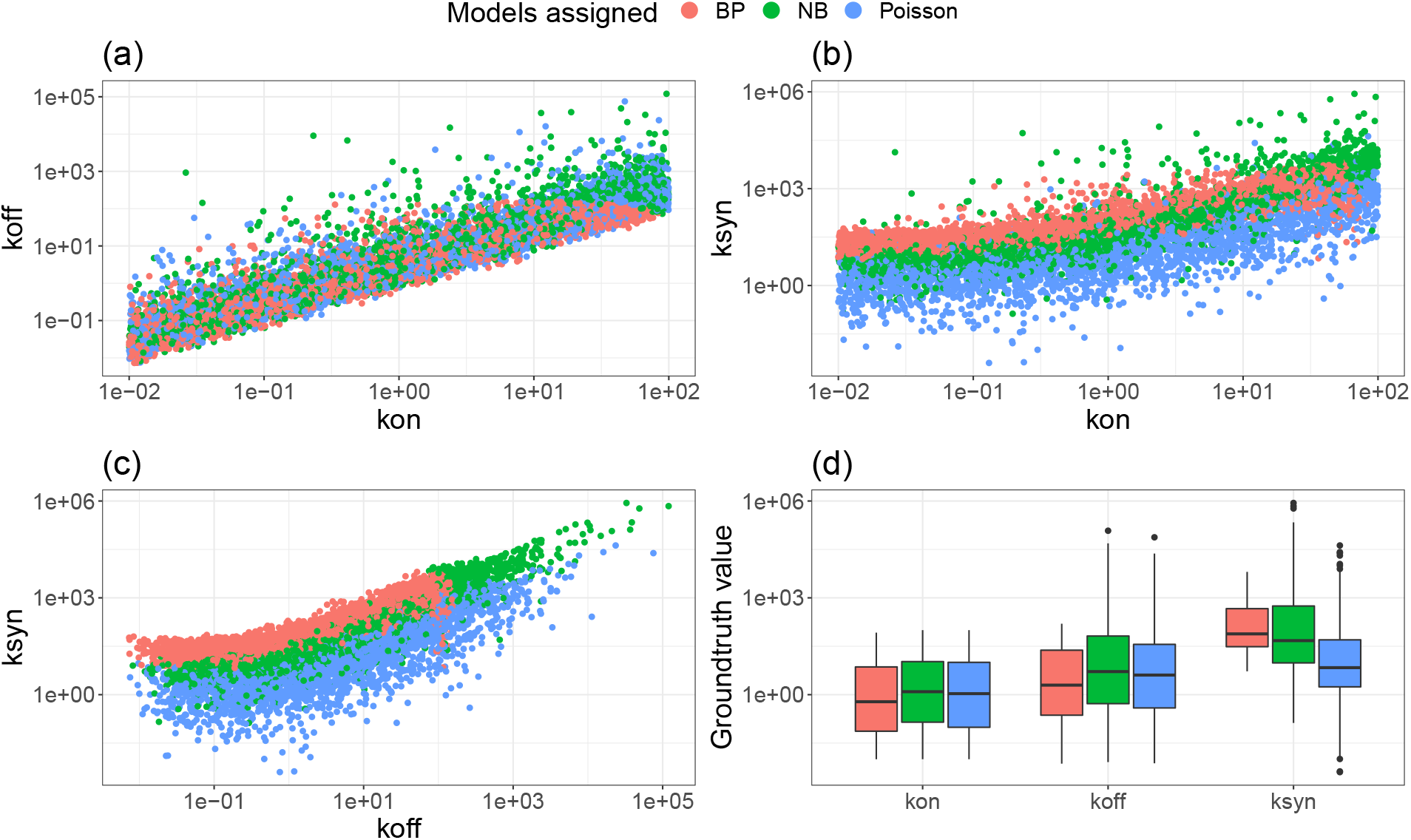
Low *k*_syn_ lead to wrong identification. **(a-c):** each dot represents one gene. Model selection of BP (Beta-Poisson), NB (negative binomial) and Poisson based on the lowest AIC. **(d)** Impact of magnitude of kinetic parameters on model selection.

**Figure S11:**
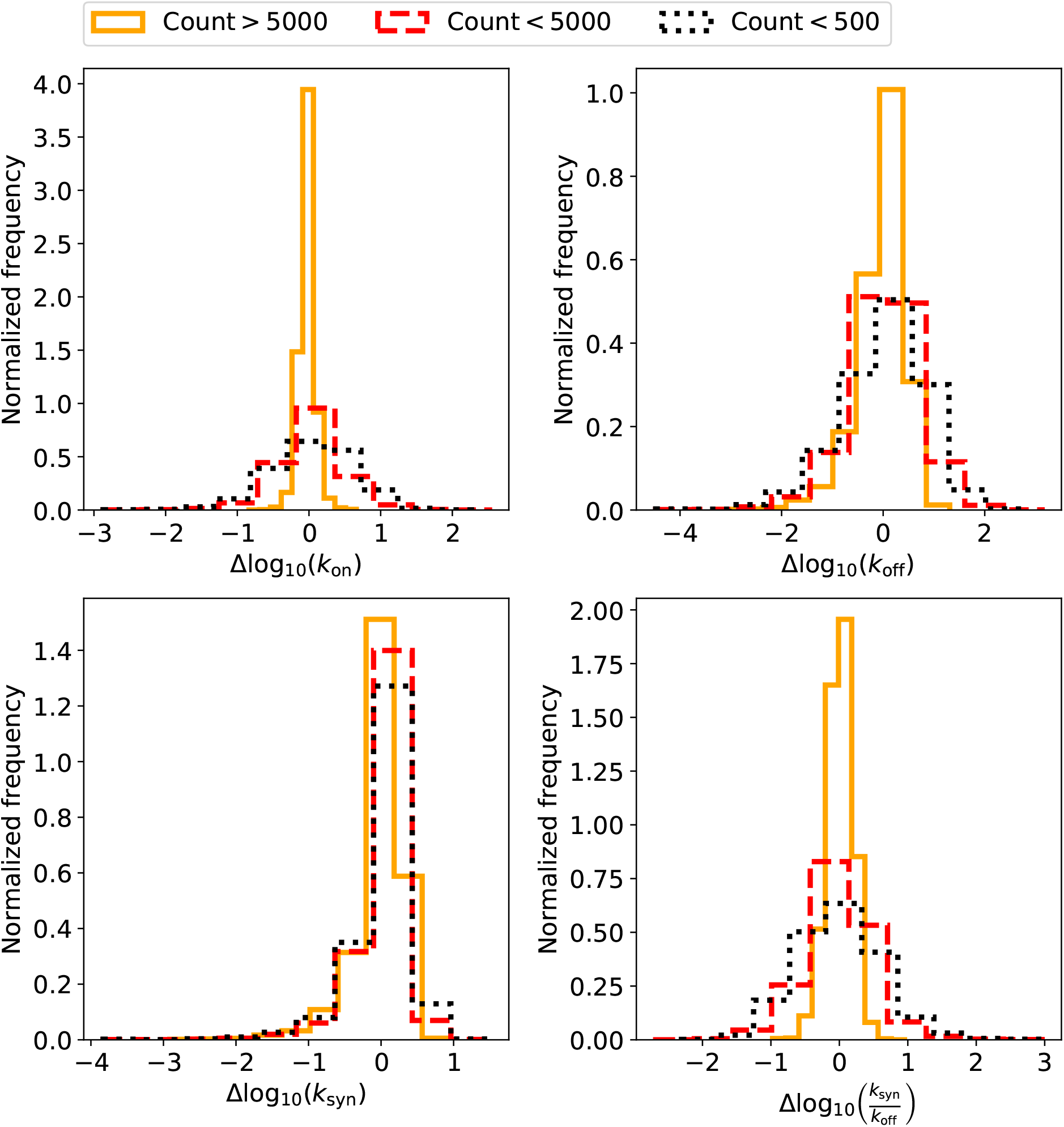
Histogram summarizing the errors in the predicted kinematic parameters for our synthetic data (Fig. S5) based on the NN for 5000 cells. The genes have been categorized based on their expression. We thus distinguish between those genes that have fewer than 500 counts, those that have fewer than 5000 counts and those that have more than 5000 counts across all 5000 cells. As can be seen from the figure, lowly expressed genes generally lead to higher errors.

**Figure S12:**
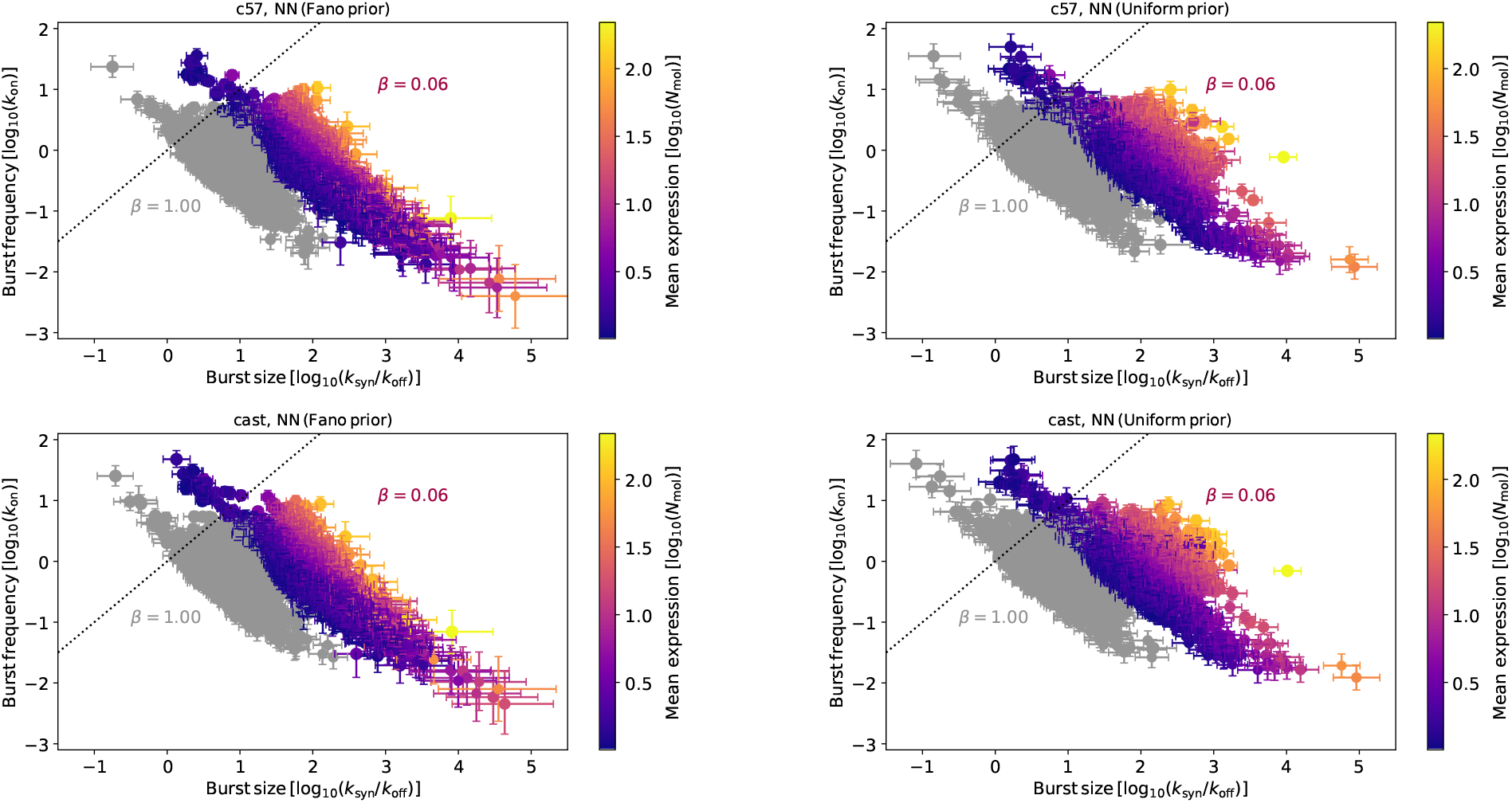
Predicted burst frequency and burst size for each allele (CAST/EiJ × C57BL/6J) based on data from [42]. The burst kinetics were inferred using the neural network. During the training, we either imposed a uniform prior for the logarithm of the Fano factor (left-hand side) or directly for the logarithm of *k*_syn_ (right-hand side). Two scenarios for the capture efficiency (*β*) were considered: A fixed capture efficiency of *β* = 1.0 (grey markers) and a varying capture efficiency with 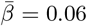. The error bars signify 68 % credibility intervals.

**Figure S13:**
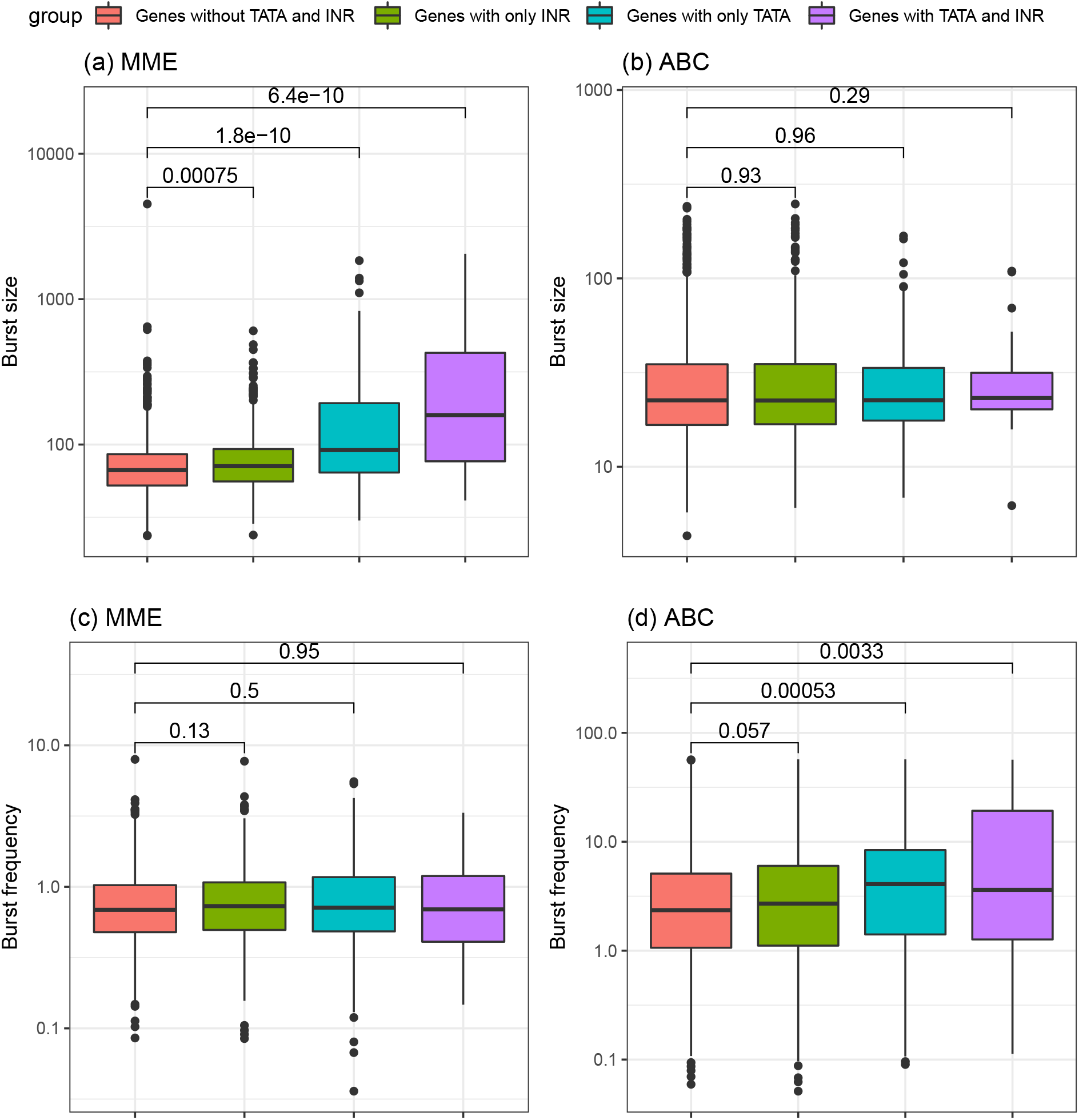
Figure corresponds to Fig. 4. Allele c57. **(a-b):** burst size inferred from MME and ABC; **(c-d):** burst frequency inferred from MME and ABC. The P-values of the Wilcox test between groups are shown.

**Figure S14:**
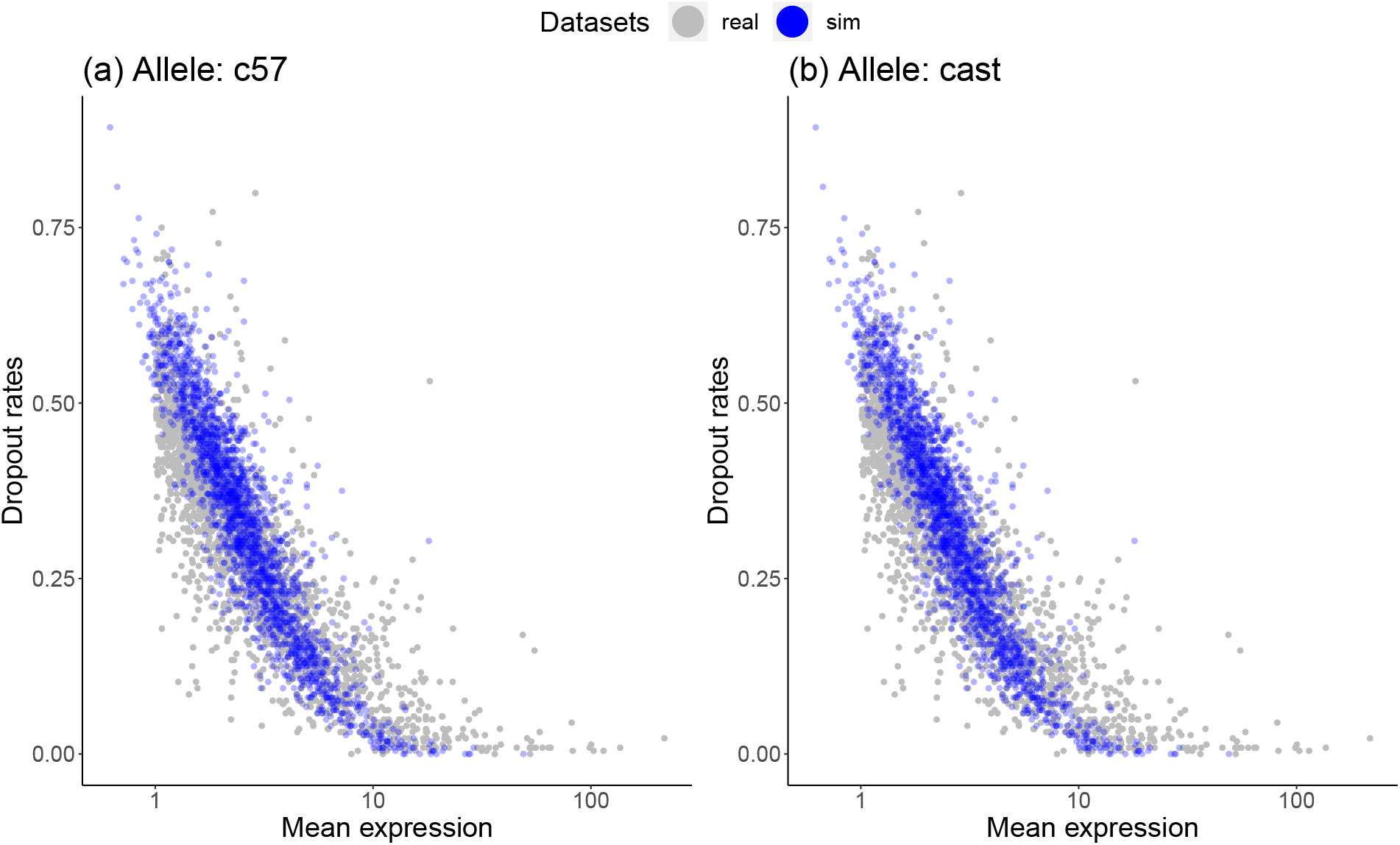
Comparison between real and simulated data in terms of the dropout-mean relationship. Simulated data was generated based on parameters estimated using the NN method. Kinetic parameters were inferred based on data from **(a)** allele c57 and **(b)** allele cast respectively.

**Figure S15:**
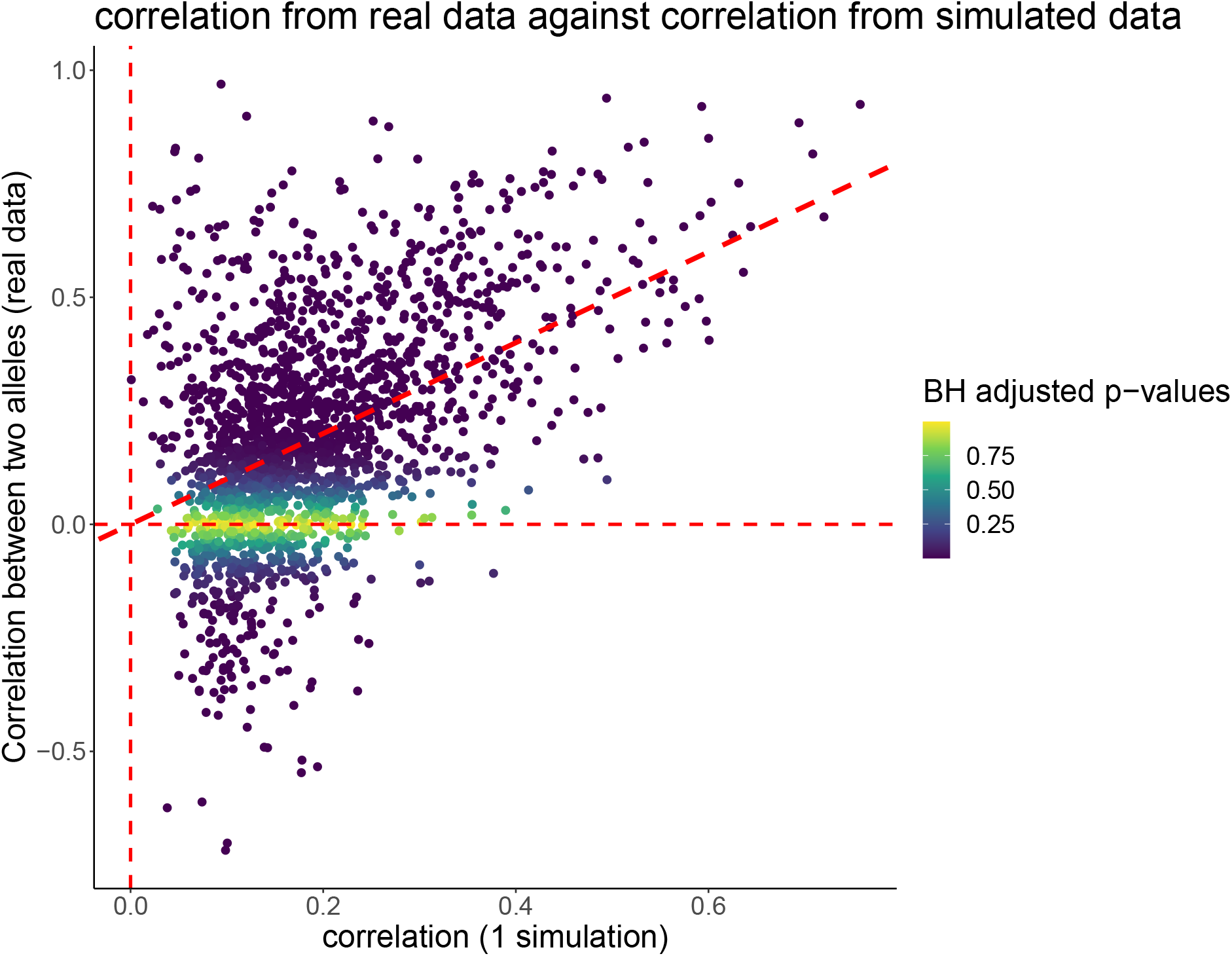
Kinetic parameters inferred from allele-specific data Larsson et al. [42]. Each point represents the Spearman correlation between two alleles for one gene. Dots are coloured according to the adjusted P-values calculated using the real data. Three dashed lines represent diagonal line; vertical lines x=0 and y=0 respectively.

**Figure S16:**
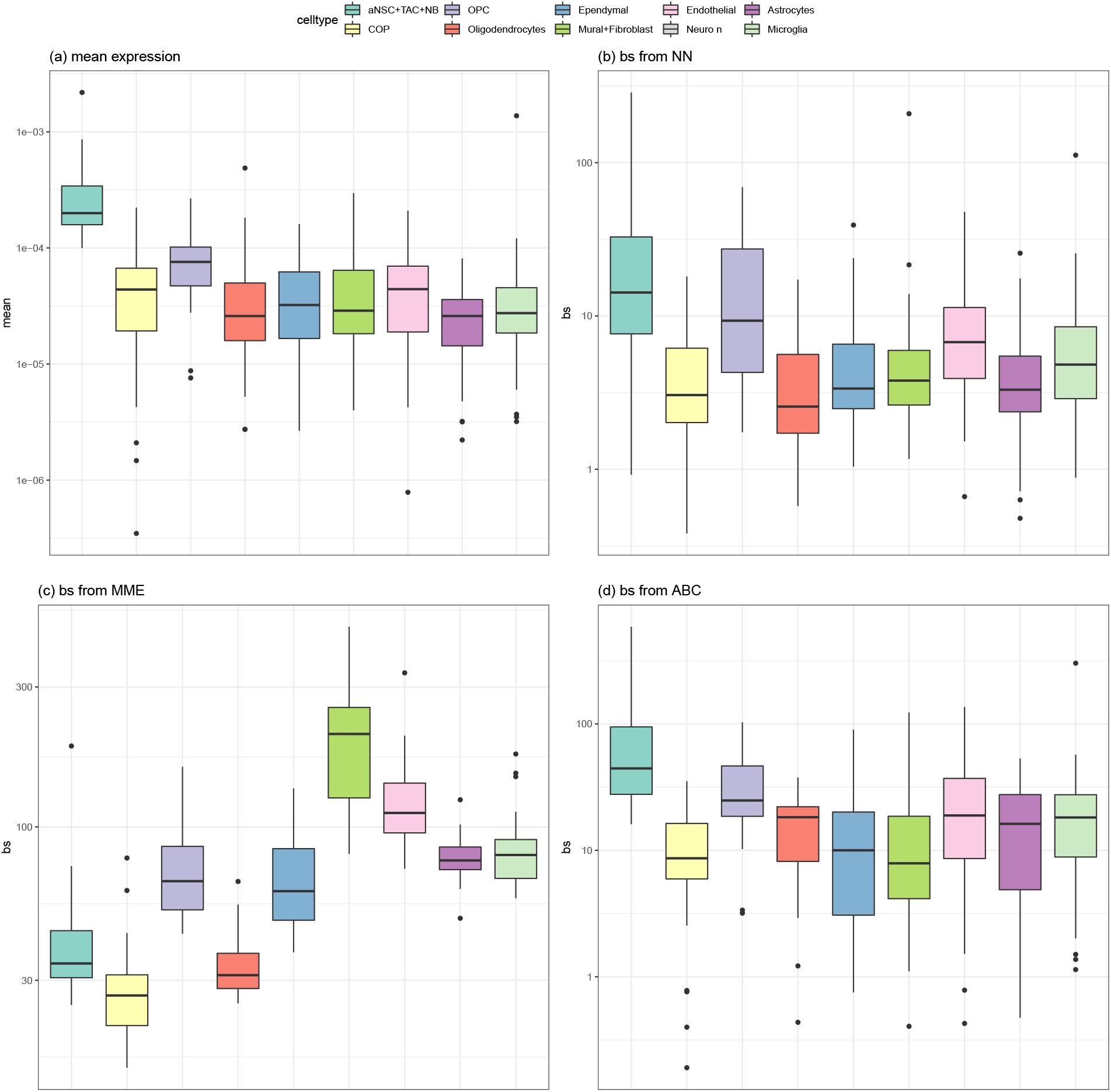
Figure corresponds to Fig. 5. **(a)** Mean expression, calculated after total count normalized; Burst size estimated using NN **(b)**, MME **(c)** and ABC **(d)**. Use aNSC marker genes reported in Mizrak et al. [58].

**Figure S17:**
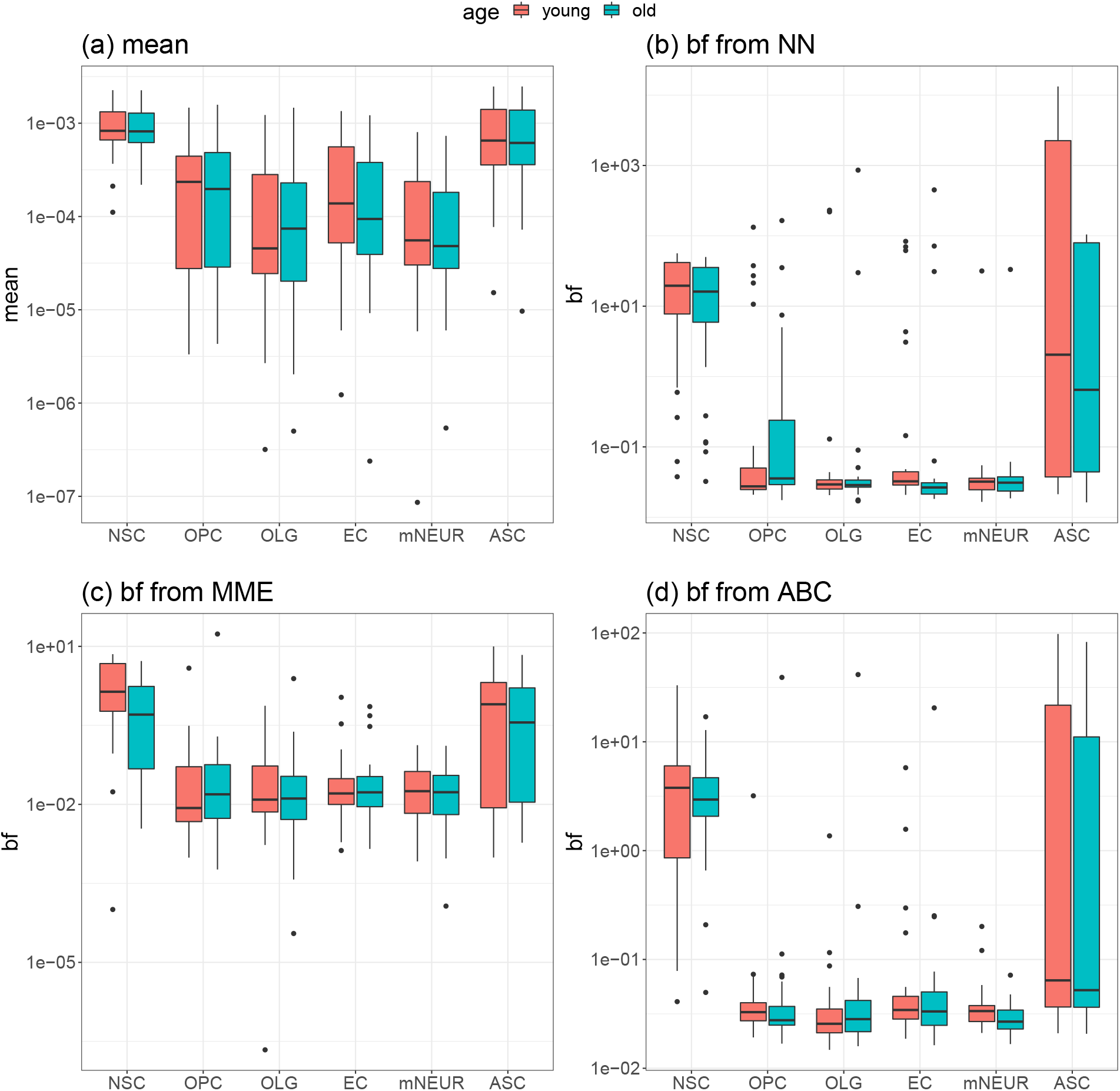
Figure corresponds to Fig. 6. **(a)** Mean expression, calculated after total count normalized; Burst frequency estimated using NN **(b)**, MME **(c)** and ABC **(d)**. Use NSC marker genes reported in Ximerakis et al. [59].

**Figure S18:**
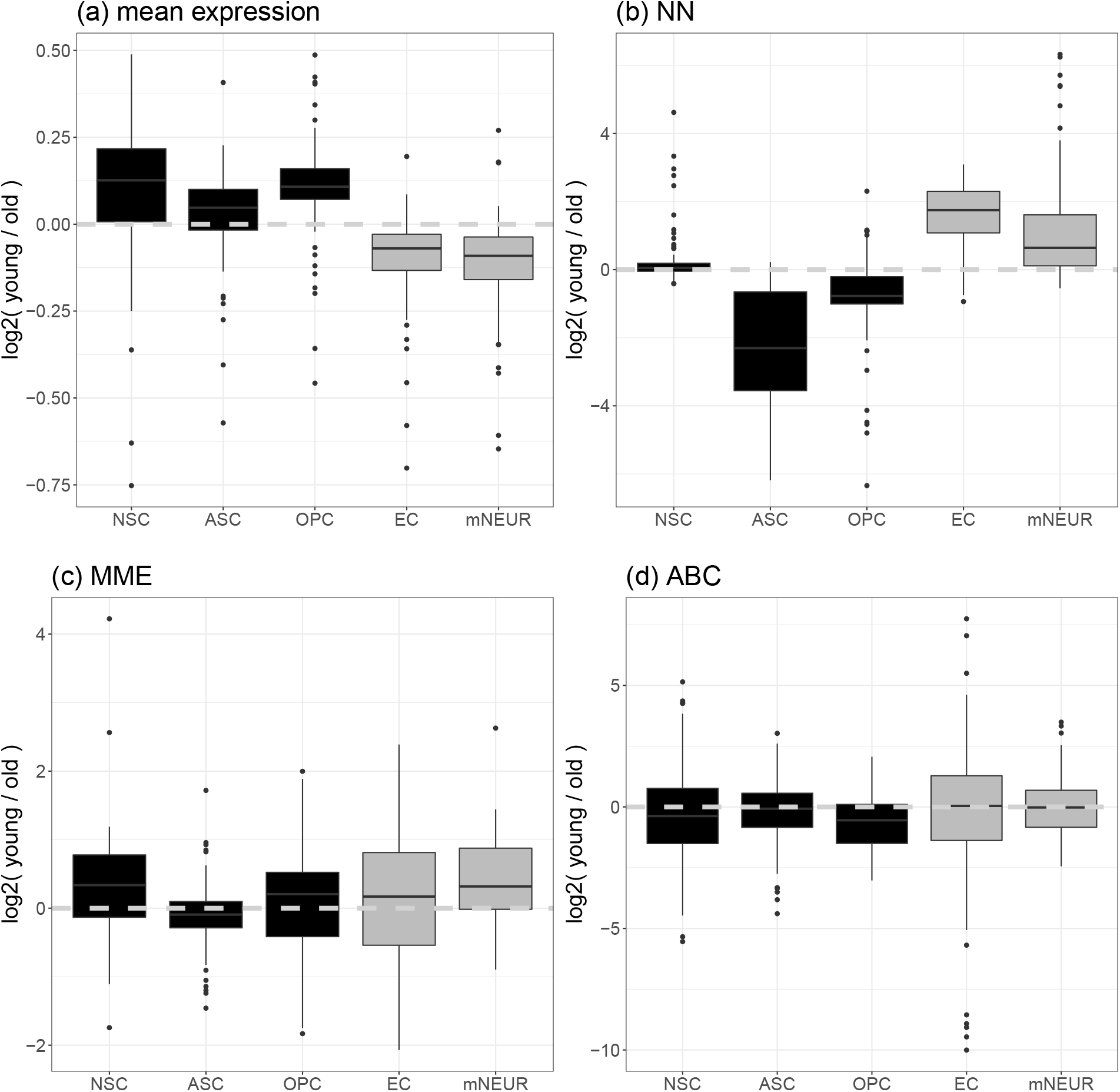
Figure corresponds to Fig. 6. Ratio of burst size of genes encoding ribosomal subunits within each cell type from young and old mice respectively. **(a)** Mean expression, calculated after total count normalized; Burst size estimated using NN **(b)**, MME **(c)** and ABC **(d)**.

